# Early human lung immune cell development and its role in epithelial cell fate

**DOI:** 10.1101/2022.12.13.519713

**Authors:** Josephine L. Barnes, Peng He, Masahiro Yoshida, Kaylee B. Worlock, Rik G.H. Lindeboom, Chenqu Suo, J. Patrick Pett, Anna Wilbrey-Clark, Emma Dann, Lira Mamanova, Laura Richardson, Amanda J. Oliver, Adam Pennycuick, Jessica Allen-Hyttinen, Iván T. Herczeg, Robert E. Hynds, Vitor H. Teixeira, Muzlifah Haniffa, Kyungtae Lim, Dawei Sun, Emma L. Rawlins, Krzysztof Polanski, Paul A. Lyons, John C. Marioni, Zewen Kelvin Tuong, Menna R. Clatworthy, James L. Reading, Sam M. Janes, Sarah A. Teichmann, Kerstin B. Meyer, Marko Z. Nikolić

**Affiliations:** UCL Respiratory, Division of Medicine, University College London, London, WC1E 6JF, UK; Wellcome Sanger Institute, Wellcome Genome Campus, Cambridge, CB10 1SA, UK; University College London Hospitals NHS Foundation Trust, London, UK; European Molecular Biology Laboratory, European Bioinformatics Institute (EMBL-EBI), Wellcome Genome Campus, Cambridge, UK; Wellcome Trust/CRUK Gurdon Institute, and Department of Physiology, Development and Neuroscience University of Cambridge, Cambridge, CB2 1QN, UK; Division of Respiratory Diseases, Department of Internal Medicine, Jikei University School of Medicine, Tokyo, Japan; Department of Medicine, University of Cambridge, Cambridge Biomedical Campus, UK; Biosciences Institute, Newcastle University, Newcastle upon Tyne, NE2 4HH, UK; Dept Physics/Cavendish Laboratory, JJ Thomson Ave, University of Cambridge, Cambridge, CB3 0HE, UK; Cambridge Institute of Therapeutic Immunology and Infectious Disease, Jeffrey Cheah Biomedical Centre, Cambridge Biomedical Campus, Cambridge, UK; CRUK Cambridge Institute, University of Cambridge, Cambridge, UK; Tumour Immunodynamics and Interception Laboratory, Cancer Institute, University College London, London, UK; Epithelial Cell Biology in ENT Research (EpiCENTR) Group, Developmental Biology and Cancer Department, Great Ormond Street UCL Institute of Child Health, University College London, London, UK; CRUK Lung Cancer Centre of Excellence, London, UK

## Abstract

During human development, lungs develop their roles of gas exchange and barrier function. Recent single cell studies have focused on epithelial and mesenchymal cell types, but much less is known about the developing lung immune cells, although the airways are a major site of mucosal immunity after birth. An open question is whether tissue-resident immune cells play a role in shaping the tissue as it develops *in utero*. In order to address this, we profiled lung immune cells using scRNAseq, smFISH and immunohistochemistry. At the embryonic stage, we observed an early wave of innate immune cells, including ILCs, NK, myeloid cells and lineage progenitors. By the canalicular stage, we detected naive T lymphocytes high in cytotoxicity genes, and mature B lymphocytes, including B1 cells. Our analysis suggests that fetal lungs provide a niche for full B cell maturation. Given the abundance of immune cells, we investigated their possible effect on epithelial maturation and found that IL-1β drives epithelial progenitor exit from self-renewal and differentiation to basal cells *in vitro*. *In vivo*, IL-1β-producing myeloid cells were found adjacent to epithelial tips, suggesting that immune cells may direct the developing lung epithelium.

## Introduction

A comprehensive understanding of human lung development on a cellular and molecular level is required to facilitate the discovery of novel therapeutic strategies for establishing effective lung regeneration and repair^1,2^. The ability to regenerate human lung tissue in the future may offer an alternative to lung transplantation to the many patients suffering with end-stage respiratory failure, the third highest cause of non-communicable disease deaths worldwide^3^. Studying fetal lung development should also enable us to improve the treatment of lung conditions that are known to be a major cause of mortality and morbidity in premature neonates^4^. Importantly, enhanced expertise in human lung development may be utilized in the development of therapies, for example in the production of patient-derived induced pluripotent stem cells (iPSCs) used for disease modeling and drug screening^5,6^.

The five overlapping stages of human lung development are well characterized morphologically^1,7,8^. For our studies, we had access to human lungs aged between 5 and 22 post-conception weeks (pcw), which includes the first three stages of lung development. In the first, embryonic stage, spanning approximately 4-7 pcw, the primary left and right lung buds appear and undergo rapid branching, forming a lobular structure. The pseudoglandular stage occurs between 5-17 pcw and consists of further growth by branching morphogenesis and establishment of the airway tree, with the appearance of smooth muscle, cartilage and mucosal glands. Blood vessels develop alongside airways, but branch more slowly than the epithelium^9^. The canalicular phase spans 16-26 pcw and includes an estimated three further rounds of epithelial branching. In addition, the airways increase in size, the distal epithelial tubes widen into airspaces and the surrounding mesenchyme thins, creating the future alveoli. Alveolar epithelial cell (AEC) differentiation begins at this stage. Our work did not cover the penultimate saccular stage (24-38 pcw) of development and the final alveolar stage (36 pcw to approximately 21 years^10^). In these final stages, saccules form distally in airways, eventually becoming capillary-wrapped alveoli via saccule septation, while AEC2s (AEC type II cells) start producing surfactant^11–15^.

In this study, we focused specifically on investigating the immune cell repertoire within developing lungs. Adult human lungs contain a plethora of immune cells (20% proportionally^16^), known to play a crucial role in both normal lung homeostasis and pathogenesis, suggestive of potential analogous functions during development. Fetal hematopoiesis begins extremely early during human development, with immune progenitors emerging from the yolk sac and aorta gonad mesonephros of embryos as early as 1-2 pcw^17,18^. These cells seed the early hematopoietic organs, the fetal liver and bone marrow^17–19^, which, by 8-9 pcw, go on to seed the lymphoid organs (thymus and spleen) and peripheral non-lymphoid organs such as the skin, kidney and gut^18,20^. Recently, it has been confirmed that immune cells are observed in human lungs as early as 5 pcw^21^.

Previous studies have demonstrated a role for immune cells in directing homeostasis^22,23^ (intestine, testis), regeneration^24^ (adult lung) and tissue development, although the latter has only been shown in the developing mouse (e.g. mammalian gland^25^). To explore the establishment of the immune system and its possible role in directing lung development, we profiled the immune cells present in early human lungs from 5 to 22 pcw, using a combination of single cell RNA sequencing (scRNAseq), CITE-seq (Cellular Indexing of Transcriptomes and Epitopes by Sequencing), B cell receptor (BCR) and T cell receptor (TCR) sequencing, immunohistochemistry (IHC) and flow cytometry. Furthermore, we functionally investigated potential immune cell-epithelial interactions *in vitro*, using three-dimensional (3D) lung epithelial organoid cultures derived from human embryonic lungs^6^ and demonstrated that immune cell-derived IL-1β can direct epithelial differentiation. We validated the expression of relevant molecules *in vivo*, using techniques including smFISH (small molecule fluorescence *in situ* hybridization) and IHC.

## Results

### Immune cells are abundant in human fetal lungs

In this study we examined developing prenatal human lungs (5-22 pcw), profiling immune cells at both the cellular and molecular level (**Fig 1A** for experimental overview). Our single cell sequencing data is initially depicted as broadly defined immune cell types (**Fig 1B**; further detail in **Fig 2**). Innate and progenitor cell types were prevalent early in development, including innate lymphoid (ILC), natural killer (NK) and myeloid cells. Over developmental time, the proportion of lymphocytes gradually increased. Given the diversity of immune cells, we also investigated the effect of immune cell signaling on epithelial differentiation as outlined in the experimental overview (**Fig 1A**).

**Figure 1.**
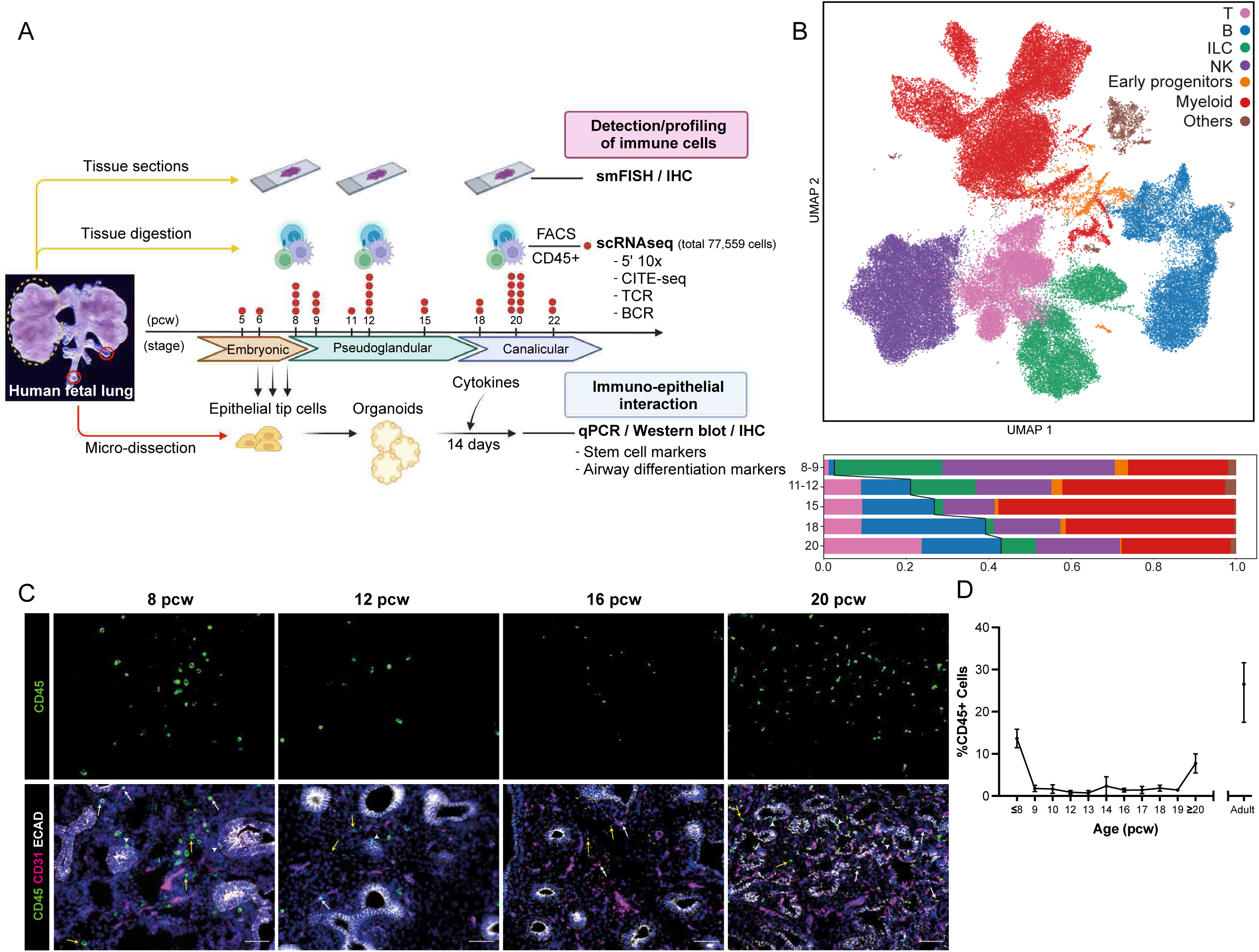
Immune cells are abundant in human fetal lungs. (**A**) Experimental overview created with BioRender.com. Human fetal lung tissue was digested and FACS-sorted to isolate CD45^+^ immune cells for scRNAseq (red dot = biological replicate; n = 31; FACS: **Fig S1A**). Tissue sections across developmental stages were used for cell type validation (IHC and smFISH), while embryonic tissue was used to generate organoids for functional studies. (**B**) UMAP (upper) and bar chart (lower) colored by broad cell populations in the single cell dataset. Representative IHC images (**C**) show the spatial distribution of CD45^+^ immune cells within the endothelium (CD31^+^, white arrows), epithelium (ECAD^+^, arrowheads) and mesenchyme (yellow arrows) during fetal lung development (blue: DAPI^+^ nuclei; scale bar = 50µM). The proportion of immune cells, as a percentage of all DAPI^+^ nuclei, was quantified in cryosections at weekly time points throughout lung development (**D**), using *ImageJ* software. Data are presented as mean ± SEM, n≥3 biological replicates.

**Figure 2.**
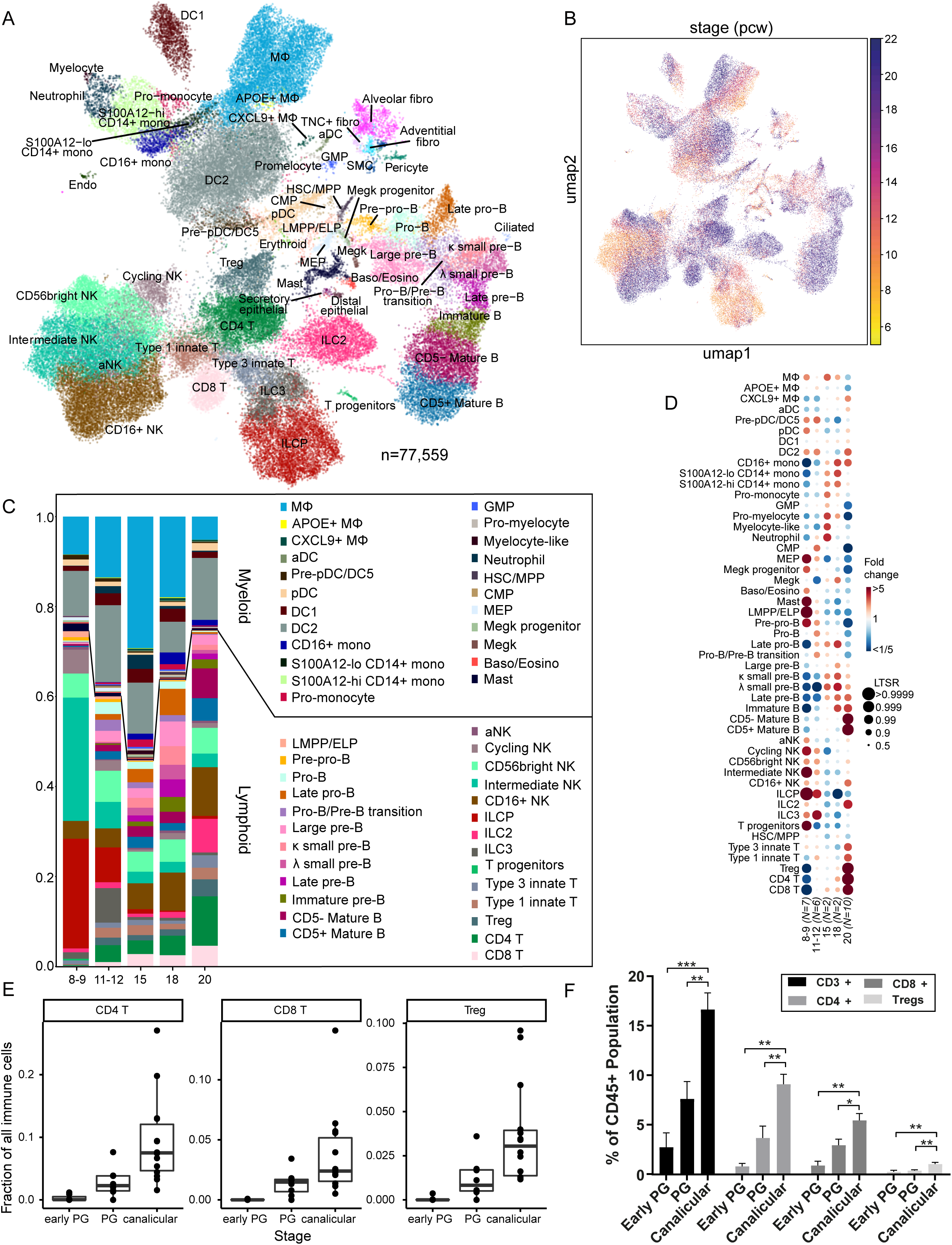
Single cell analysis of fetal lung immune cells. (**A**) Single-cell transcriptome profiles embedded onto a 2D UMAP plane, colored by cell type/state and (**B**) sample developmental age. (**C**) Proportions of each cluster across age groups. (**D**) Dot plot showing fold change in the proportions of lung-resident immune cell types across fetal age, estimated by fitting a generalized linear mixed model. Each dot is color-coded by the fold change over the mean of each donor group, and scaled by its probability using the local true sign rate (LTSR). (**E**) Barplot showing the abundance of CD4, CD8 and Treg cells throughout lung development, as a fraction of all scRNAseq immune cells. (**F**) Flow cytometric analysis (gating: **Fig S1B**) of digested whole fetal lungs shows the proportions of CD3^+^ T cells at stages throughout lung development, separated into: the whole T cell population (CD3^+^), CD4^+^ cells, CD8^+^ cells and Tregs, calculated as a proportion of the entire CD45^+^ immune cell population. Early PG = 7-9 pcw, PG = 10-14 pcw and Canalicular = 17-21 pcw (‘PG’ = pseudoglandular). Data are presented as mean ± SEM, n=3 biological replicates. See also **Fig S2, S3** and **Table S1**.

IHC demonstrated that CD45^+^ immune cells were abundant in human fetal lungs throughout early development **(Fig 1C**) and were located within all lung regions, including the endothelium (CD31^+^, incorporating the vasculature), epithelium (E-Cadherin^+^ (ECAD^+^)) and mesenchyme. Our images showed that CD45^+^ immune cells were prevalent in lung tissue up to 8 pcw, followed by a relative decrease, before increasing again after 20 pcw. We quantified IHC in whole lung tissue sections at weekly time points from 5 to 22 pcw, in a minimum of three replicates at each (**Fig 1D**), and confirmed the expression pattern observed in the representative IHC images, revealing a peak in the proportion of immune cells in the early pseudoglandular stage up to 8 pcw, which subsequently declined; the proportion of immune cells then increased again after 20 pcw, at the canalicular stage. As expected, adult lungs contained the same proportion of immune cells as previously reported (~20%)^16^, approximately double the maximum number we observed in the fetus at the half-way point of normal gestation. We conclude that there exists a complex and dynamically changing immune compartment in the fetal lung.

### Molecular characterization of immune cells using scRNAseq

To comprehensively characterize the subpopulations of fetal lung immune cells, we digested whole fetal lungs aged 8, 9, 12 and 20 pcw and enriched CD45^+^ immune cells by FACS (**Fig S1A**). These cells were profiled using 10X Chromium 5’ scRNAseq with a subset also used for CITE-seq protein measurement, with quality control metrics given in **Fig S2A,B**. To maximize cell type and cell-state discovery, we combined these data with the immune compartment of our previous unbiased single-cell whole fetal lung scRNAseq data^21^. After filtering out cells with fewer than 200 genes detected, BBKNN^26^ integration was performed, followed by downstream clustering and dimensionality reduction (see **Methods**). After careful curation and removal of cell clusters with high mitochondrial reads, low gene numbers and doublet signatures, we obtained a total of 77,559 high quality (**Fig 2A**) transcriptomic profiles, covering all known leukocyte lineages, including B and T lymphocytes, innate lymphoid cells (ILCs), natural killer (NK) cells, myeloid cells and the early progenitors of the aforementioned lineages (**Fig 1B**). Non-immune cells (“others”) such as epithelial, erythroid and stromal cells were also identified and are likely to represent cells non-specifically bound by CD45 antibodies. Although not immune cells, we included these cells in our data object.

We carefully annotated and curated the clusters of the single-cell transcriptomic profiles across all stages, presenting 59 clusters of cell types/states (**Fig 2A**) based on marker gene expression (**Fig S2C**) as described in existing literature (**Table S1**). Early progenitor cells, including the pre-pro-B cell, the lymphoid-primed multipotent progenitor cell (LMPP), innate lymphoid cell progenitor (ILCP), megakaryocyte-erythroid progenitor (MEP), common myeloid progenitor (CMP), megakaryocyte progenitor and T cell progenitors are mainly present in the earlier stages, as visualized in the stage-specific UMAP and cell type proportion bar and dot plots (**Fig 2B,C,D**). To quantitate changes in cell type proportion, we used a mixed linear model to calculate the fold-change of cell type proportions over time (**Fig 2D**). This confirmed the overrepresentation of progenitors, in particular ILCPs and LMPPs/ELPs (early lymphoid progenitors), T progenitors and MEPs at 8-9 pcw. However, a number of non-progenitor cell types, such as macrophages, mast cells and NK cells are also overrepresented early in development, in line with reports that innate immune cells develop prior to the establishment of the adaptive immune system^18^. Over developmental time, we also see a gradual increase in monocytes. For the T cell lineage we detect some early progenitors, whilst single positive T cells are first observed at 12 pcw with no detection of intermediates. Their proportions are highly upregulated at 20 pcw. In contrast to T cells, B lymphocyte development occurs gradually. The developmental trajectories for B lymphocytes, NK/T and innate lymphoid cells are examined in more detail in the next sections. Finally, myeloid lineage cells, including macrophages, dendritic cells, megakaryocytes and granulocytes are present from early stages.

To validate our scRNAseq results (**Fig 2D,E**), we carried out flow cytometry, which confirmed the significant increase in abundance of total T cells (CD3^+^), CD4^+^, CD8^+^ and regulatory T cells with age (**Fig 2F, S1B**). At the early pseudoglandular stage, proportions of CD4^+^ and CD8^+^ cells were negligible, as expected based on our knowledge of human thymic development^27^. The proportions significantly increased during the pseudoglandular and canalicular stages, with a CD4/CD8 ratio (1.25 and 1.67, respectively) in range of that considered normal in the healthy adult lung^28^. The proportion of Tregs within the CD4^+^ population was between 10% and 15%, and remained relatively stable across each stage of development. We then confirmed the presence of identified immune cell types by IHC in tissue sections and observed positive staining for the T cell markers CD4 and CD8, for the NK cell marker CD56 and the B lymphocyte marker CD20 at both 12 and 20 pcw (**Fig S3A**). We validated progenitors and rare cell types using RNAscope (**Fig S3B,C**), including extremely rare SMIM24^+^SPINK2^+^ cells, likely to be HSCs as indicated by the expression pattern of these markers (**Fig S3D**). In fact, we observed a continuous spectrum of developing myeloid lineages *in silico* and spatially validated GMPs & CMPs, promonocytes & myelocytes, MEPs & megakaryocyte progenitors and megakaryocytes in 18 and 20 pcw fetal lung samples, based on their canonical marker gene expression (**Fig S3C,D**). This suggests that myeloid cells can exit the fetal liver at premature stages and continue to differentiate in the lung.

We see a wealth of immune cell types within the developing lung. Notably the total fraction of immune cells is high at the embryonic stage at 8 pcw and again at the canalicular stage of 20 pcw (**Fig 1D**). We speculated that the early peak was partially due to the presence of maternal immune cells which are known to enter the fetus^29^. To investigate such events in our single cell dataset, we used genetic deconvolution (see **Methods**), which has been used successfully to detect maternal cells in the embryonic placenta^30^, and found that the proportion of maternal cells was negligible. It is conceivable that vascular maturation may contribute to the increase in immune cells from 20 pcw, to account for the second observed peak (**Fig 1D**). To assess the proportion of vasculature in the developing lung, we used *PECAM1* as a surrogate readout. Expression of this gene in endothelial cells increases in embryonic development, but remains relatively constant, on a per-cell basis after this period (**Fig S3E**). RT-PCR quantification showed a strong increase in *PECAM1* expression from the embryonic to pseudoglandular to canalicular stage (**Fig S3F**). Given its relatively constant expression per cell from 12 pcw onwards, this analysis suggests that the proportion of vascular cells does increase at the canalicular stage. Hence, the dynamic increase in immune cell number coincides with the increasing mass of vasculature. In conclusion, we propose that the peaks of immune cells are likely to reflect consecutive waves of immune populations egressing from primary lymphoid organs, with innate immune cells being more prevalent in the early phase and mature B and T lymphocytes becoming abundant later.

### A B cell developmental niche in human fetal lungs

While examining the B cell compartment, we were able to detect all developmental stages of B cells (**Fig 3A-C, Table S1**) in prenatal lungs, including LMPP/ELP (*CD34^+^EBF1^−^*), pre-pro-B (*EBF1^+^SPINK2*^+^*VPREB1^+^*), pro-B (*DNTT*^+^), large pre-B (*IL7R*^+^*MS4A1*^+^*MKI67*^+^), small pre-B (*IL7R*^+^*SPIB*^+^*MKI67^−^*), immature B (*MS4A1*^+^*IGHD^lo^IGHM^hi^VPREB3^hi^*) and mature B (*MS4A1*^+^*IGHD^hi^IGHM^hi^VPREB3^lo^*) cells. We were able to define more precise cell subtypes based on cluster-specific genes: *NEIL1*^+^*MKI67^−^* late pro-B cells, *RAG1*^+^*MS4A1^+^* pro-B/pre-B transition cells, small pre-B cell clusters, expressing either immunoglobulin kappa (*IGKC*^+^) or lambda (*IGLC2*^+^*IGLC3*^+^) light chain genes, *MS4A1*^+^ late pre-B cells that expressed the marker for surrogate light chain, *IGLL1*, which forms part of the pre-B cell receptor, and *CD5^−^* versus *CD5^+^* mature B cells. To validate the accuracy of annotations, we performed a label transfer using a logistic regression-based method (CellTypist^31^) with the prenatal immune cell atlas from other organs^20^ as training data, and showed that our manual annotations were overall consistent with predicted labels (**Fig S5A**, **Table S2**). Trajectory analysis was performed using monocle3, with HSC/MPP (hematopoietic stem cells/multipotent progenitor cells) set as a starting point (**Fig 3B**). The arrangement of cell types along the UMAP overall follows a linear trend and is consistent with the known biology of B cell maturation^20,32,33^, progressing from pre-pro-B to pro-B, pre-B and mature B cell stages (cell markers in **Fig 3C**). **Fig 3D** visualizes the top 100 differentially-expressed genes along this trajectory, with a subset further enlarged in **Fig S4A**. We see the acquisition of B cell markers such as *IL7R* at the pre-pro-B cell stage, persisting towards the pro-B/pre-B transition, in line with its essential role in lineage progression^34^. The trajectory recapitulates the transition from cycling (*MKI67*) to non-cycling stages, that sit between the BCR heavy and light chain rearrangement events, with expression of *DNTT* for generating junctional diversity, plus biphasic expression of recombinase activating genes *RAG1* and *RAG2* at the pro-B and pre-B cell stages. As expected, the later stages of B cell maturation coincide with the appearance of *MS4A1* (*CD20*) and *IgD* (**Fig 3D**, **S4E**). Interestingly, while *IGHM* transcription is observed from the earliest stages (**Fig S4E,F**), the surface expression increases dramatically only at the immature B cell stage for IgM and mature B cell stage for IgD.

**Figure 3.**
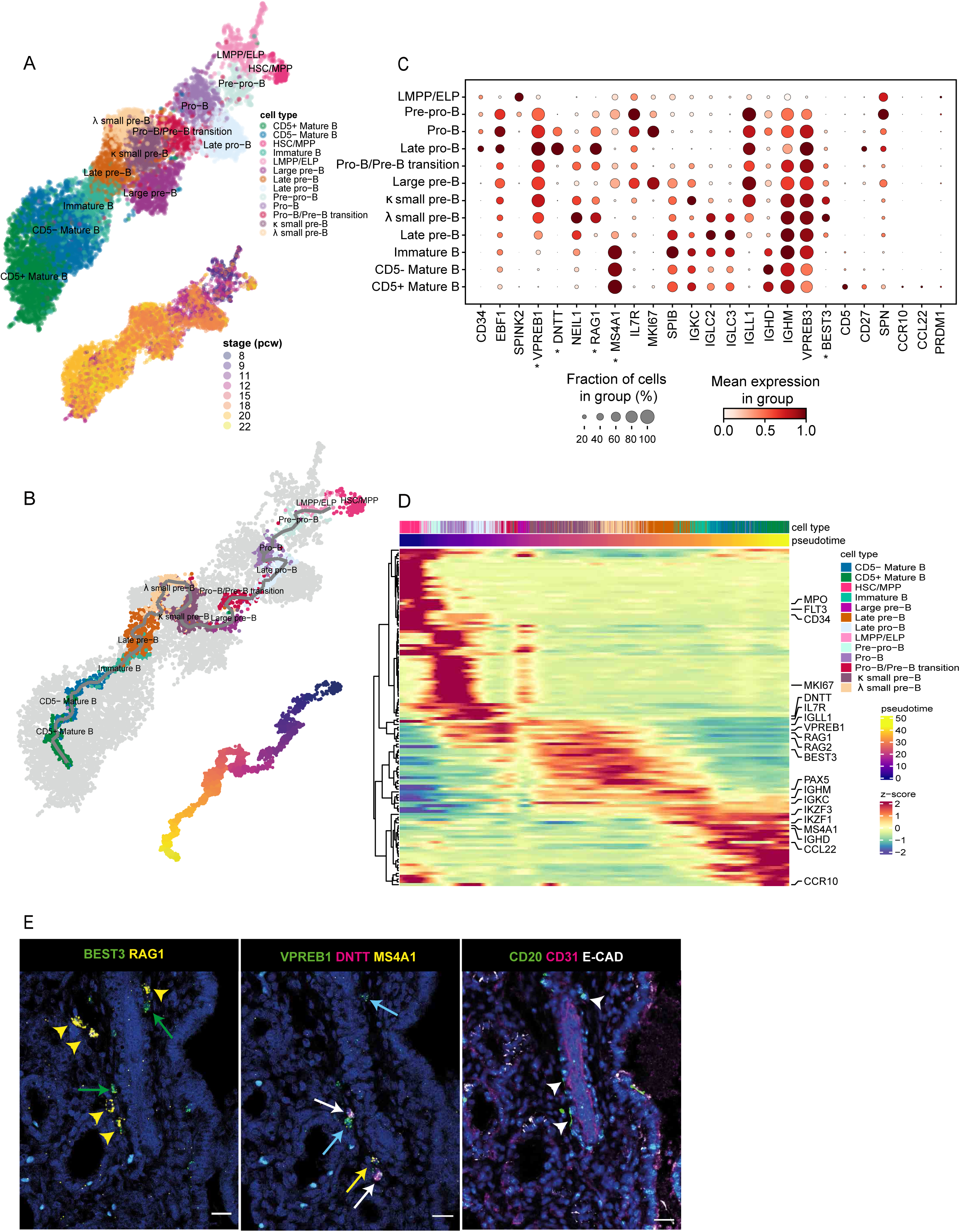
B cell development in fetal lungs. (**A, B**) UMAPs of the B cell lineage showing (**A**) cell type clusters (top) and developmental stage of the corresponding samples in post-conception weeks (bottom); and (**B**) inferred trajectory from HSC/MPP to mature B cells using monocle3 (top) and the corresponding pseudotime (bottom). (**C**) B cell marker gene expression. (**D**) Heatmap showing expression of the top 100 genes, with differential expression along the trajectory shown in (**B**). The cells (columns) are ordered by pseudotime and genes (rows) are arranged by hierarchical clustering and optimal leaf ordering. RNAscope using sequential tissue sections from 20 pcw fetal lungs (**E**, left and center) shows B cells expressing multiple combinations of B progenitor markers (labeled with asterisks in (**C**)), *BEST3*, *RAG1*, *VPREB1*, *DNTT* and *MS4A1* (yellow arrowheads (pre-B): *BEST3^+^RAG1^+^*, green arrows (large pre-B): *BEST3^+^*, white arrows (pro-B): *VPREB1^+^DNTT^+^*, blue arrows (large pre-B): *VPREB1^+^*, yellow arrows (late pre-B): *VPREB1^+^MS4A1^+^*). Corresponding IHC (**E**, right) using the next sequential tissue section, shows expression of CD20, CD31 (endothelium/blood vessels) and ECAD (epithelium) (arrowheads: CD20^+^ B cells). See also **Fig S4**.

Until recently, it was believed that B cell maturation mainly occurs in the fetal bone marrow, although a recent paper from our laboratory has demonstrated that B cell intermediates can also be detected in fetal skin, kidney and gut^20^. Here, we show that all populations representing the B cell developmental trajectory can also be found in the fetal lung. To understand whether these come from the circulation, potentially “leaking” out of the bone marrow, or whether these develop *in situ*, we performed smFISH and IHC and observed that developing B cells of different stages clustered together (**Fig 3E, S4D**). RNAscope showed *VPREB1^+^DNTT^+^* cells representing pro-B cell stages, *RAG1^+^BEST3^+^* small pre-B cells and *MS4A1^+^* mature B cells, while CD31, EpCAM and CD20 staining on consecutive tissue sections demonstrated that these B lineage cells map to the extravascular space (**Fig 3E, S4D**). Taken together, our findings provide strong evidence that B cells develop locally in the fetal lung, supporting the previous notion^20^ that B lymphocyte development can occur in the periphery, outside of primary hematopoietic organs.

### Putative B-1 cells in fetal lung

To further characterize the mature B cell populations in the fetal lung, we examined Ig isotype expression and clonal expansion. Both RNA and CITE-seq protein measurements (**Fig S4E**) show that the large majority of mature CD5^+^ and CD5^−^ B cells express IgM and IgD, with only a small fraction of cells having switched to IgG or IgA isotypes. Absence of the expression of the transcription factor *PRDM1* (**Fig 3C**) confirms that our dataset does not contain any plasma cells, consistent with the notion that the fetal lung environment is largely sterile.

The final stage of mature CD5^+^ cells are marked by *CD5*, *CD27*, *SPN* (*CD43*), *CCR10* and *CCL22* (**Fig 3C**), representative of putative human B-1-like cells (**Fig S5A**) that were previously reported in human fetal bone marrow, gut, liver, kidney, skin, spleen and thymus^20^. Moreover, at the single-cell BCR repertoire level, CD5^+^ mature B cells displayed a shorter CDR3 length (heavy chain) (**Fig S4B**) than CD5^−^ mature B cells. The same pattern is demonstrated by the random non-coded/palindromic (N/P)-insertion lengths (**Fig S4C**), suggesting that DNTT/RAG activity during V(D)J rearrangement is reduced in these cells. These observations are consistent with the B-1-like cells previously reported^20^. It is possible that the developing lung epithelium and mesenchyme support the homeostasis of B-1 cells through B-1 cell-specific chemokine pathways such as CCL28-CCR10 and DPP4-CCL22 (**Fig S5C**).

### T, NK and innate lymphoid cells in fetal lungs

Further investigating other cell types of the lymphoid lineage, we identified conventional and unconventional (type 1 and 3 innate) T cells, ILCs and NK cells (**Fig 2A** and **4A**), plus a tiny cluster of T progenitors (*PTCRA*^+^*RAG1*^+^*RAG2*^+^; **Fig S5D**) that are likely contaminants from the thymus. CD4, CD8 and Treg T cells bear the hallmarks of naive T cells (**Fig 4B**). To investigate the maturity of these T cells, we compared their transcriptional identity to naive T cells encountered after birth. To this end, we integrated fetal lung CD4 and CD8 T cells with naive T cells derived from PBMCs (peripheral blood mononuclear cells) from neonates and pediatric donors^35^ and performed differential expression analysis over age. We observed a prominent age-associated transcriptional signature in naive CD4 and CD8 T cells during fetal stages and in early childhood, revealing unique characteristics of fetal naive T cells (**Fig S5F**). Strikingly, genes that are more highly expressed in fetal life include cytotoxicity markers such as *GZMA* and *NKG7*. This could suggest that naive T cells exhibit an innate immune function during fetal development, which is lost after exposure to pathogens during childhood. Interestingly, fetal-specific naive T cell genes remain expressed up until the end of neonate age, indicating that naive T cells continue to mature even after birth.

**Figure 4.**
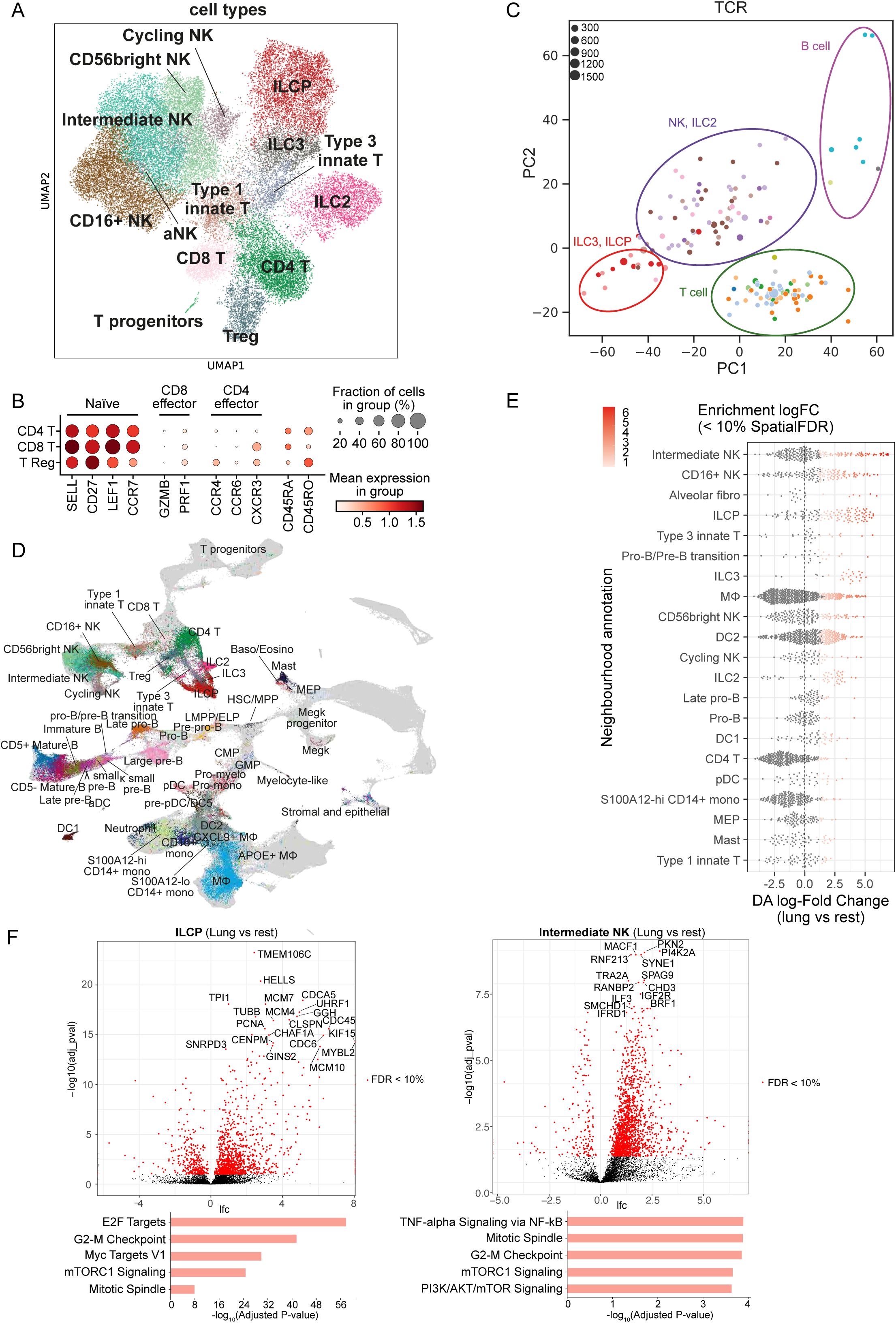
T cells, ILCs and NK cells in fetal lungs. (**A**) UMAP showing lymphocytes except B cells. (**B**) Expression of marker genes for naive and mature T cells. (**C**) Principal Component Analysis (PCA) plot summarizing TRBJ and TRBC gene segment usage proportion in different cell types. Each dot represents a sample of at least 20 cells, with dot size representing the cell count. Colored circles illustrate groupings of the cell types. (**D**) UMAP embedding scVI integrated fetal immune cells from lung and 9 hematopoietic, lymphoid and non lymphoid tissues. Fetal lung cells are colored by their cell type annotation. (**E**) Beeswarm plot showing the distribution of log fold change in abundance between lung cells and all other organs in neighborhoods containing cells from different lung cell type clusters (using annotations from cross-tissue fetal immune atlas^20^). Only differential abundance neighborhoods at SpatialFDR 10% and logFC > 0 are colored. (**F**) Differential gene expression comparing fetal lung with the other organs in ILCP and intermediate NK cells. Below are the top 5 enriched biological processes GO terms for upregulated genes. See also **Fig S5 and S6**.

Like previous observations in other developing organs^20^, we see an abundance of unconventional T cells *in situ* in developing lung, but the equivalent is not apparent in adult lung (**Fig S5B**). A cluster of ILCPs were identified by expression of marker genes (*HPN* and *SCN1B*)^36^; **Fig S5E**). One important question is whether they are local progenitors of all ILC subtypes or of a specific subtype. Interestingly, 20-40% of ILC and NK cells expressed nonproductive TCRꞵ chain with fewer proportions expressing nonproductive TCRɑ, ɣ, or δ chains (**Fig S6A-D**), and the non-productive TCRꞵ chains were mainly contigs, consisting of the J segment with the C segment, but no V region (**Fig S6E**). This is consistent with previous reports of murine ILCs having undergone partial TCR recombination^37,38^. Next, we investigated the J and C segment usage pattern in TCRꞵ for different cell types (**Fig S6G,H**) and summarized the repertoire grouped by sample with principal component analysis (PCA; **Fig 4C**). On the PCA, ILCPs co-localized with ILC3s, while ILC2s mainly co-localized with NK subtypes. Thus ILCPs share V gene segment usage with ILC3s. As ILCPs do not express *RAG* (**Fig S5D**), no further VDJ recombination happens, and J/C usage in nonproductive TCRꞵ should be preserved in local differentiation. This suggests that, in the lung, ILCPs can potentially only give rise to ILC3s but not ILC2s.

### Higher levels of ILCPs in fetal lung

To further explore our data, we integrated our fetal lung immune population with immune cells from other developing organs^20^ using scVI (**Fig 4D**), allowing us to search for cell neighborhoods enriched in the developing lung (**Fig 4E**). ILCs and NK cells proved to be significantly enriched in the fetal lung compared to other fetal tissues at similar developmental stages. Lung ILCPs and intermediate NK cells differentially expressed genes associated with positive regulation of the cell cycle compared to other fetal organs, suggesting that these progenitor/differentiation intermediates are in a proliferative state and that the fetal lung may provide a niche to expand these lineages (**Fig 4F**). Although ILC2s are known to be present in multiple fetal organs and directly contribute to the pool of adult ILC2s^39^, it is possible that a higher baseline of fetal lung ILC2s is particularly needed to prepare for the rapid activation of an IL-33/IL-13-driven immune response upon birth, which regulates lung homeostasis and tolerance^40^.

### IL-1β causes tip stem cells to exit from a self-renewing state and differentiate to basal cells in fetal lung organoids

In human developing lungs, the distal epithelial tips are composed of SOX9^+^ progenitors, which give rise to all alveolar and airway lineages^6^. Having established that immune cells were prevalent throughout the developing lung, we wondered which immune cells were specifically located around the epithelial tips at each stage during development, which cytokines they may be secreting and the subsequent effect they may have on epithelial maturation, specifically on tip progenitors. IHC showed CD45^+^ immune cells adjacent to SOX9^+^ epithelial tips in fetal lungs (**Fig 5A**), with more immune cells clustered around tips at 8-9 and 20 pcw than at 12 pcw (**Fig 5B**), which may reflect the biphasic peaks that we observed previously at these time points (**Fig 1C,D**). To study immune-epithelial interactions we used an established organoid model, whereby distal epithelial tips are microdissected from fetal lung tissue aged 5-9 pcw and cultured to form self-renewing, branching, three-dimensional (3D) organoids^6^. We used previously published bulk RNAseq data, to compare cytokine receptor expression in organoids derived from 5-9 pcw lung epithelial tips with freshly-dissected 6-7 pcw tips^6^ (**Fig 5C**). This showed that epithelial tip progenitors express several cytokine receptors, including receptors to IL-1, IL-4, IL-6, IL-13, IL-17, IL-22, IFN-γ, TNF and TGF-β. We found analogous expression in fresh epithelial tips and organoids in a specific subset of cytokine receptors. Based on the receptors expressed, we screened the effects of the corresponding cytokines on cultured organoids.

**Figure 5.**
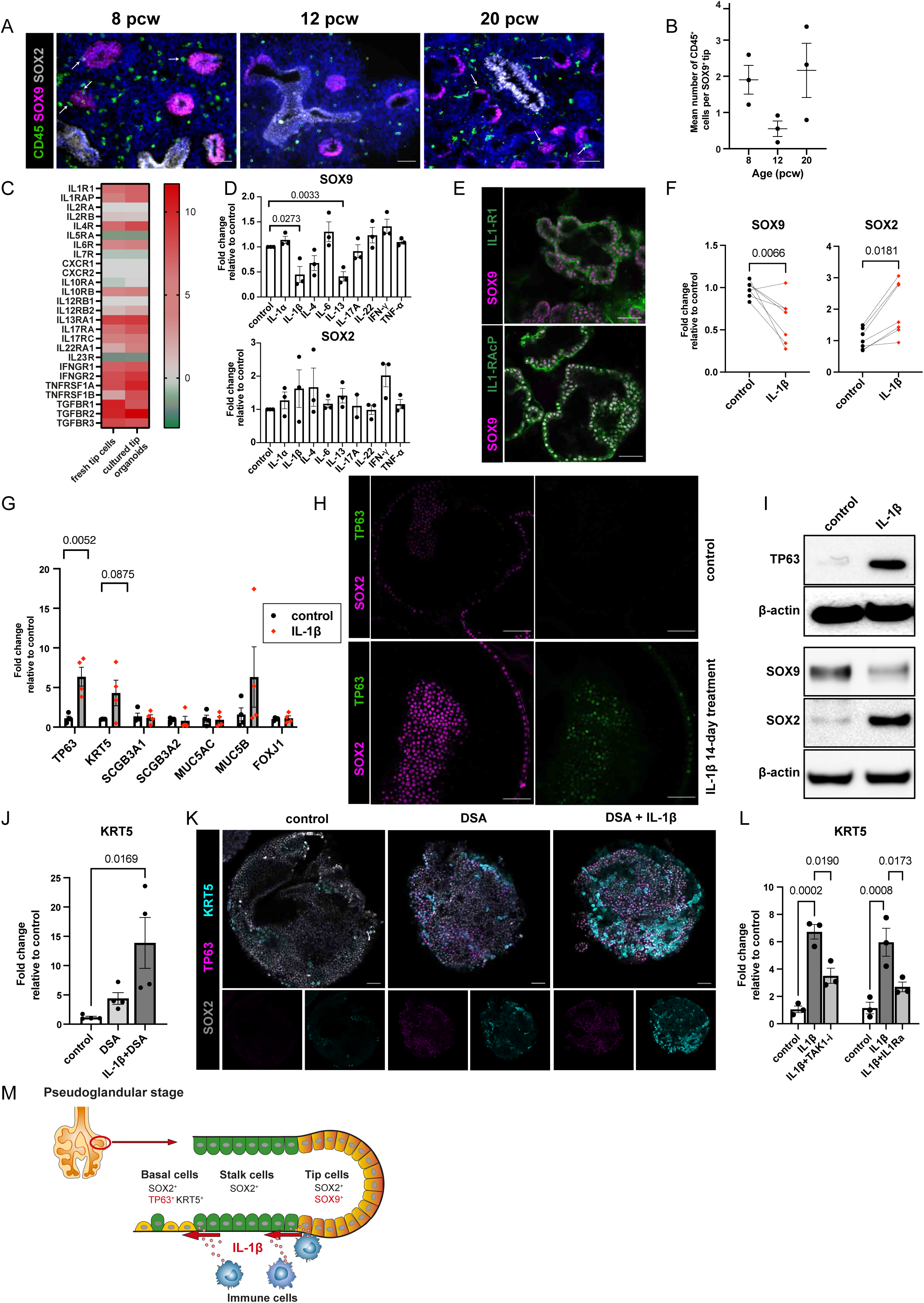
IL-1β causes exit from a self-renewing state and differentiation to basal cells in fetal lung organoids. (**A**) Representative IHC images show the spatial distribution of CD45^+^ immune cells surrounding SOX9^+^ epithelial tips during lung development (white arrows: immune cells in direct contact with SOX9^+^ cells; blue: DAPI^+^ nuclei; scale bar = 50µm). (**B**) IHC images were quantified at 8-9, 12 and 20 pcw to determine the number of immune cells making direct contact with SOX9^+^ tip cells. (**C**) Heatmap showing a comparison of cytokine receptor expression in RNAseq of fresh epithelial fetal lung tips versus cultured organoids^6^. (**D**) RT-PCR of *SOX9* and *SOX2* in organoids treated with listed cytokines for 7 days. (**E**) Whole mount staining shows lung organoid IL-1R1 and IL-1R1AcP expression (scale bar = 50µm). (**F**) *SOX9* and *SOX2* expression analysis by RT-PCR after IL-1β stimulation for 14 days. (**G**) RT-PCR analysis of differentiation markers: *TP63* and *KRT5* (basal cell markers), *SCGB3A1*, *SCGB3A2, MUC5AC* and *MUC5B* (secretory cell markers), and *FOXJ*1 (ciliated cell marker). (**H**) Whole mount staining shows expression of SOX2 and TP63 in lung organoids treated with or without IL-1β for 14 days (**H**; scale bar = 50µm). (**I**) Western blotting shows SOX9, SOX2 and TP63 expression in organoids cultured with or without IL-1β for 14 days (β-actin = loading control). (**J**) The effect of IL-1β on basal cell differentiation after treating organoids for 3 days with TGF-β and BMP4 (dual-SMAD activation/DSA) alone or combined with IL-1β, assayed by RT-PCR measuring *KRT5*. (**K**) Whole mount organoid staining shows SOX2, TP63 and KRT5 expression following DSA/IL-1β-treatment (scale bar = 50µm). (**L**) Organoid differentiation was tested via RT-PCR of KRT5 after treatment with two IL-1 signaling inhibitors, TAK1 inhibitor (TAKi), and an IL-1R antagonist (IL-1Ra), alone and in combination with IL-1β. (**M**) Model of the effect of IL-1β on nearby lung epithelial tips, whereby IL-1β causes a decrease in SOX9 expression, with exit from a self-renewing state, and an increase in SOX2, TP63 and KRT5, marking differentiation to basal cells. All data are presented as mean ± SEM, n≥3 biological replicates; treated samples are normalized to untreated control samples; p-values are calculated by one-way ANOVA followed by Tukey’s post-hoc test (**B, J, L**) or unpaired two-tailed t test (**D, F, G**). See also **Fig S7**.

Distal epithelial tip cells co-express SOX9 and SOX2 throughout the pseudoglandular stage (~5-16 pcw)^5,6,41^. As development proceeds, SOX9 disappears, while SOX2 increases and cells move proximally, to become SOX9^−^/SOX2^+^ airway progenitors^6,42^. Since fetal lung organoids are long-term self-renewing cultures derived from embryonic/early pseudoglandular distal epithelial tip progenitors, they maintain co-expression of SOX2 and SOX9 throughout culture^6^ (**Fig S7B**). Alterations in their expression are, therefore, a useful measure of response to stimuli and hence we utilized SOX2 and SOX9 as markers for realtime PCR analysis, investigating the effects of our cytokine panel. Initially, organoids were treated with 10 ng/ml cytokines for 7 days (**Fig 5D**) and we observed that IL-1β and IL-13 caused a significant decrease in *SOX9* expression, while *SOX2* was not significantly altered. The decrease in *SOX9* caused by IL-13 may be explained by its known role in promoting goblet cell differentiation, which has been observed in the human adult lung^43^, but the IL-1β result was not expected. Therefore, we decided to further explore the effect of IL-1β signaling on organoids.

First, we checked for the presence of the IL-1 receptor (IL-1R1) on the surface of organoids. For IL-1β signal transduction, both IL-1R1 and an accessory protein (IL-1RAcP) are required. Whole mount staining showed both IL-1R1 and IL-1RAcP expression in organoids (**Fig 5E**), confirming the feasibility of IL-1β signal transduction. Importantly, the expression of IL-1R1 and IL-1RAcP were also confirmed in fetal lung epithelium (**Fig S7C,D**). Knowing that both receptor components were present, we cultured organoids in the presence of IL-1β for a longer time period. After 14 days of IL-1β treatment, we observed a significant decrease in *SOX9* and increase in *SOX2* (**Fig 5F**). Therefore, the effect observed following organoid-treatment with IL-1β, suggests it causes the withdrawal of tip progenitors from a self-renewing state towards differentiation. We hence proceeded to look at the effect of IL-1β treatment (10ng/ml, 14 days) on several markers of airway differentiation, and found that it caused a significant increase in the expression of *TP63* and a non-significant increase in *KRT5*, both markers of basal cells; but it had no effect on the expression of secretory cell markers *SCGB3A1*, *SCGB3A2*, *MUC5AC*, *MUC5B* or the ciliated cell marker *FOXJ1* (**Fig 5G**). The effect of IL-1β treatment on organoids was also confirmed at the protein level, via whole mount staining (**Fig 5H**) and Western blotting (**Fig 5I**), which showed that IL-1β caused a decrease in SOX9 and an increase in SOX2 and TP63. These results suggest that IL-1β promotes the exit of tip progenitor cells from a self-renewing state by downregulating SOX9 and increasing SOX2 expression and then causing an increase in TP63 expression, thereby promoting differentiation to basal cells.

Using an equivalent fetal lung organoid model, it has previously been shown that dual SMAD signaling activation (DSA) via TGF-β and BMP-4 induces differentiation of lung epithelial tip progenitors into immature TP63^+^ basal cells^44^. We found that IL-1β supplementation in conjunction with DSA caused a significant increase in the mature basal cell marker, KRT5, in lung organoids (**Fig 5J,K**). This suggests that immune cells that secrete IL-1β promote mature basal cell differentiation and may work in conjunction with mesenchymal cells, which secrete TGF-β/BMP-4^44^. To further confirm the role of IL-1 signaling in differentiation, we also looked at the effect of IL-1β inhibition. IL-1 receptor blockade via IL-1 receptor antagonist (IL-1Ra) increased proliferation (Ki67^+^ cells) and organoid size. Conversely, IL-1β decreased organoid size and proliferation (**Fig S7E,F,G**). This suggests that IL-1 signaling plays an important role in determining progenitor cell fate, either in stemness or differentiation. Furthermore, adding IL-1Ra plus IL-1β significantly attenuated IL-1β-induced basal cell differentiation (**Fig 5L**). The MAPKKK protein kinase, transforming growth factor β-activated kinase 1 (TAK1), mediates activation of JNK and NF-κB in the IL-1-activated signaling pathway^45,46^. (5Z)-7-Oxozeaenol is a potent inhibitor of TAK1 and was also used to treat organoids for 14 days to inhibit IL-1β-mediated signaling. TAK1 inhibition significantly attenuated IL-1β-induced basal cell marker induction, further confirming the key role of IL-1 signaling (**Fig 5L**).

### Myeloid cells secrete IL-1β in fetal lungs

Having established that IL-1β signaling has the capacity to change epithelial cell fate in the developing lung, we wanted to identify those immune cells that may be a potential source *in vivo*. The primary sources of IL-1β in adults are blood monocytes, tissue macrophages, and dendritic cells^47,48^. We first confirmed the existence of *IL1B^+^* immune cells *in vivo* in fetal lungs using RNAscope (**Fig 6A**). In our single cell dataset, the top 5 highest expressing cell types were the source of more than 75% of all *IL1B* (**Fig 6B,C**). Fetal DC2s, macrophages, monocytes and neutrophils showed the highest *IL1B* expression (**Fig 6B,C, Fig S8A**), *IL1RN* (**Fig 8B**) and *CASP1* expression (**Fig S8C**). The expression of *IL1B* in the epithelium itself in the pseudoglandular stage was undetectable (**Fig S7H**), further supporting myeloid cells as the main source of IL-1β in early lung development. We investigated the spatial and temporal distribution of dendritic cells and macrophages in the developing lungs more closely, utilizing CD1C as a marker for DC2 dendritic cells, and CD206, the macrophage mannose receptor, as a marker for macrophages (**Fig 6D,E**). We identified CD1C^+^ DC2 dendritic cells and CD206^+^ macrophages in fetal lung tissue throughout development and we found that there were dendritic cells/macrophages directly interacting with SOX9^+^ epithelial tip progenitors. In addition, we also observed monocytes and neutrophils near the developing epithelium (**Fig 6H**). We hypothesize, therefore, that several myeloid cells, including DC2s, macrophages, monocytes and neutrophils located adjacent or near to SOX9^+^ epithelial tip progenitors secrete IL-1β, initiating IL-1β signal transduction in those tip cells. This promotes exit from a self-renewing state by first downregulating SOX9 and increasing SOX2 expression, and subsequently increasing TP63 and KRT5 expression, leading to differentiation into basal cells (**Fig 5M**).

**Figure 6.**
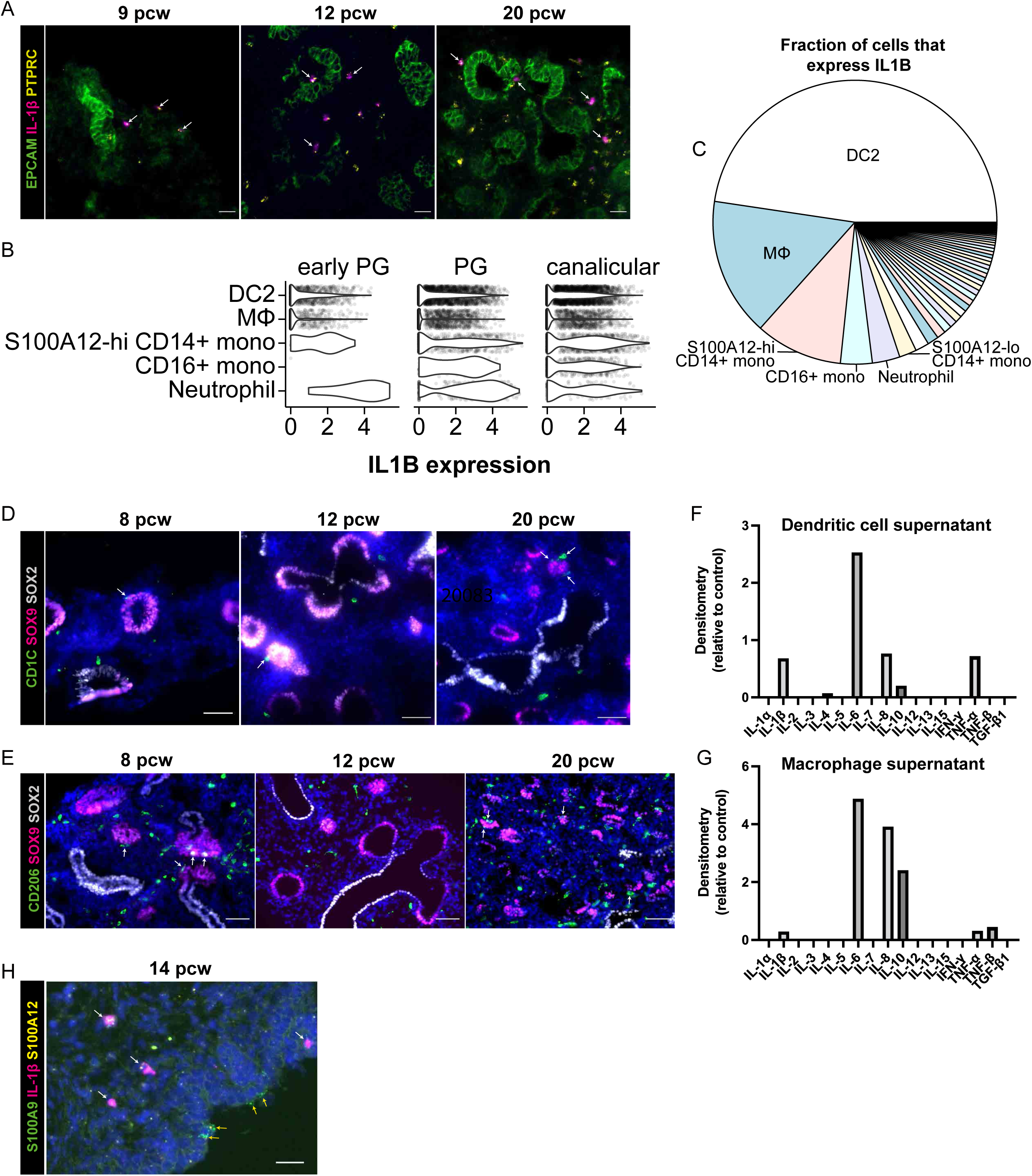
Fetal lung myeloid cells secrete IL-1β. (**A**) Representative RNAscope images of fetal lungs aged 9, 12 and 20 pcw, showing expression of *PTPRC* and *IL1B*, with EpCAM IHC (white arrows: *PTPRC^+^IL1B^+^* cells; scale bar = 20µM). (**B**) Violin plot showing *IL1B* gene expression in each of the top 5 highest expressing cell types, based on our single cell dataset (see **Fig S8** for all cell types). (**C**) Pie chart showing the total contribution of each cell type to all expressed *IL1B* mRNA. Representative IHC images show the distribution of CD1C^+^ DC2 dendritic cells (**D**) or CD206^+^ macrophages (**E**) surrounding SOX9^+^ epithelial tips during lung development (white arrows: cells adjacent to SOX9^+^ cells; blue: DAPI^+^ nuclei; scale bar = 50µM). Isolated dendritic cells or macrophages (via FACS of 21 pcw lungs, **Fig S1C**) were cultured for 7 days to investigate cytokine production. The pooled supernatant, from days 3, 5 and 7 of culture, was analyzed using the Human Cytokine Antibody Array (abcam; **F** and **G** respectively, n=1). (**H**) Representative RNAscope image showing the distribution of *S100A9^+^S100A12^+^* neutrophils/monocytes relative to the epithelium (determined morphologically), including those that coexpress *IL1B* (white arrows) and those that do not (yellow arrows) (blue: DAPI^+^ nuclei; scale bar = 20µM).

We wanted to investigate the properties of fetal lung myeloid cells in more depth, to confirm or refute our hypothesis, so we isolated dendritic cells and macrophages from fetal lungs (21 pcw). To investigate their cytokine secretion, each cell type was cultured alone for 7 days, using optimized conditions. The supernatant was removed (and replaced) on day 3, 5 and 7, pooled and analyzed using the Human Cytokine Antibody Array (abcam) (**Fig 6F,G**). Several cytokines were secreted by both cell types, notably IL-1β, IL-8, IL-6, IL-10, TNF-α and TNF-β. Importantly, this data confirms that the dendritic cells and macrophages present in developing fetal lungs are able to secrete IL-1β. IL-6 and TNF-α were originally tested via organoid-treatment for 7 days (**Fig 5D**) and had no effect on *SOX9* or *SOX2* expression. IL-8 signals via the CXCR1 and CXCR2 receptors, which are both absent from epithelial tips (**Fig 5C**), and hence IL-8 signaling may not directly affect the developing epithelium. IL-10 signaling requires both components of the IL-10 receptor (α and β), but only IL-10Rβ is expressed in epithelial progenitors and organoids (**Fig 5C**), suggesting that IL-10 also cannot directly affect epithelial tips. Although IL-8 and IL-10 cannot directly signal to the epithelium, it is possible that they may influence the recruitment and activation of other immune cells that are present in the developing fetal lung, and hence could indirectly affect the epithelium.

Overall, we have shown that immune cells capable of secreting IL-1β are resident in the vicinity of the epithelial tips. This suggests that immune cells have the capacity to direct the developing lung epithelium.

## Discussion

During embryogenesis, tissue resident immune cells become established throughout the body, laying the foundations for immune surveillance after birth and throughout life. In addition, immune cells may perform functions in tissue remodeling^49–52^ and organogenesis^53,54^, yet the prenatal immune cell repertoire has only been profiled in a handful of organs. Here we have profiled the immune cell compartment of fetal lungs from 5 to 22 pcw. We identified 59 different cell types and states, including very immature hematopoietic precursors, restricted lineage progenitors as well as more mature myeloid and lymphoid cells. Given the ability of immune cells to secrete cytokines, we examined and demonstrated the ability of cytokines to affect epithelial differentiation, pointing towards a possible broader role of the immune cells in coordinating organogenesis.

Our data suggest that the fetal lung microenvironment supports the full gamut of B lymphocyte differentiation, whilst T cells are present in the lung at the single positive stage from 12 pcw, with increased numbers at 20 pcw. Whilst there is evidence of the presence of a microbiome in the human fetal lung as early as 9 pcw^55^, the vast majority of lymphocytes we detect are naive, with little evidence of T cell activation, B cell class switching or plasma cell differentiation. This suggests that any interactions with a potentially existing microbiome are rare, or do not lead to cell activation, or that a mostly sterile environment prevails.

Lung development is mediated through the complex interaction of multiple cell types. Recent human prenatal studies have largely focused on epithelial and mesenchymal interactions, with none examining the role of immune cells. For example, recent analysis demonstrated that dual SMAD signaling activation mediated through mesenchymal cells robustly increased TP63 expression in human lung epithelial tip organoids; surprisingly this did not affect SOX9 expression^56^. During the pseudoglandular stage of development, cell proliferation is highest at the distal tip of growing buds, which co-express SOX9 and SOX2. As these tip cells differentiate into the airway lineage, they initially downregulate SOX9 expression to become more proximal stalk cells^6^. A recent CRISPRi screen experiment has demonstrated that the transcription factor SOX9 maintains epithelial tip progenitor function by promoting proliferation and inhibiting airway differentiation^57^. However, the molecular mechanism of SOX9 downregulation in airway progenitors remained unknown. Here, we demonstrate that IL-1β decreases SOX9 expression and proliferation, resulting in airway differentiation. The converse effect of IL-1β and IL-1Ra on tip proliferation further supports the concept that IL-1 signaling can affect epithelial progenitor cell fate decision and highlights the distinct role of early innate immune cells in the airway differentiation niche.

Of the cytokines released by immune cells, IL-1β has been implicated in human neonatal lung injury and in bronchopulmonary dysplasia (BPD)^58–60^. Transgenic IL-1β overexpression in mice is known to disrupt normal lung development^61^. These detrimental effects of IL-1β are pronounced during the saccular stage, but not observed in earlier stages^62^. The level of IL-1β signaling is temporally regulated and plays an important role in airway development, but over- or prolonged expression of IL-1β in later stages may inhibit alveolarization by skewed differentiation into the airway lineage.

In mouse lung development, a p63^+^ multipotent airway progenitor population emerges at E9.0 (embryonic stage), followed by p63^hi^ cells acquiring cytokeratin KRT5 to become mature tracheal basal cells at E14.5-15.5 (pseudoglandular stage)^63^. This mirrors our previous human fetal lung data^21^, which shows that basal cells mainly expand during the pseudoglandular stage of development (**Fig S7A**). Here, in our single cell dataset, we were able to demonstrate *IL1B* transcription in myeloid cells from the early pseudoglandular stage through to the canalicular stage (**Fig S8A**) and we spatially detected IL-1β-expressing immune cells in close proximity to the SOX9^+^ tip progenitors in these developmental stages (**Fig 6A,D,E,H**). These results suggest that IL-1β-secreting fetal myeloid cells stimulate the emergence of a TP63-positive population and promote further maturation of basal cells at the pseudoglandular stage (**Fig 5M**). The precise role of immune cells in the alveolar niche in later stages of development remains to be elucidated.

Whilst, we have focused our functional analyses on the role of cytokines in early epithelial development, our study charts the presence of a complex and highly dynamic immune compartment, providing an important resource for the scientific community on which to base future functional studies to examine the interplay of the immune compartment with endothelial, epithelial or mesenchymal cells.

## Materials and Methods

### Human embryonic and fetal lung tissue

Human embryonic and fetal material was provided by the Joint MRC/Wellcome Trust (grant # MR/R006237/1) Human Developmental Biology Resource (www.hdbr.org). Lung tissue was obtained from terminations of pregnancy spanning 5 to 22 pcw. Fresh lung tissue was collected in Hibernate E medium (ThermoFisher Scientific, A1247601), unless otherwise specified. Samples were staged according to their external physical appearance and measurements. Samples used had no known genetic abnormalities (karyotype analysis was undertaken on every sample by HDBR).

### Human adult lung tissue

Fresh healthy adult lung tissue (background tissue from lobectomies for lung cancer) was obtained from University College London Hospitals NHS Foundation Trust (as part of the study: An Investigation into the Molecular Pathogenesis of Lung Disease II, REC reference: 18/SC/0514, IRAS project ID: 245471).

### RNA isolation and realtime PCR for whole fetal lung

All solutions and consumables used were RNase free. Fresh lung tissue was collected in Hibernate E medium or RNAlater Stabilization reagent (ThermoFisher Scientific, AM7020). A maximum of 30 mg of tissue (or 15-20 mg if stored in RNAlater) was taken and disrupted using a micro-pestle in 600 μl of RLT buffer containing β-mercaptoethanol (10 μl/ml). Lysate was homogenized using QIAshredder spin columns (Qiagen, 79656) and total RNA was extracted with the RNeasy Mini Kit (Qiagen, 74104), according to the manufacturer’s instructions. RNA purity and concentration was determined using the Nanodrop 1000 spectrometer (ThermoFisher Scientific).

RNA was reverse transcribed using 5X qScript cDNA Supermix (Quantabio, 733-1177), according to the manufacturer’s protocol. Briefly, 0.5 μg RNA was used per sample and cDNA was diluted 1:2 with RNAse free water. For each real-time PCR reaction, 1 μl cDNA was added to 5 μl of Power SYBR Green Master Mix (Life Technologies Ltd), 0.25 - 2 μl of 10 mM forward and reverse oligonucleotide primer mix concentration optimized previously (**Table 1**), in a final volume of 10 μl in RNAse-free water. Each primer was designed using Primer3 software and synthesized to order by Sigma. For each gene, the reaction was run in triplicate and a ‘no template control’ was included. After a 10 min enzyme activation step at 95°C, 40 x PCR cycles, consisting of a 15 s denaturation step at 95°C, followed by an annealing and extension step at 60°C, were carried out. Data was collected using Quant Studio Real-Time PCR Software v1.1. Relative transcript quantities were calculated using the ΔΔCt method, normalized to the expression of GAPDH. Fold-change was calculated relative to the embryonic stage, with p-values obtained by one-way ANOVA, followed by Tukey’s post-hoc test.

**Table 1.**
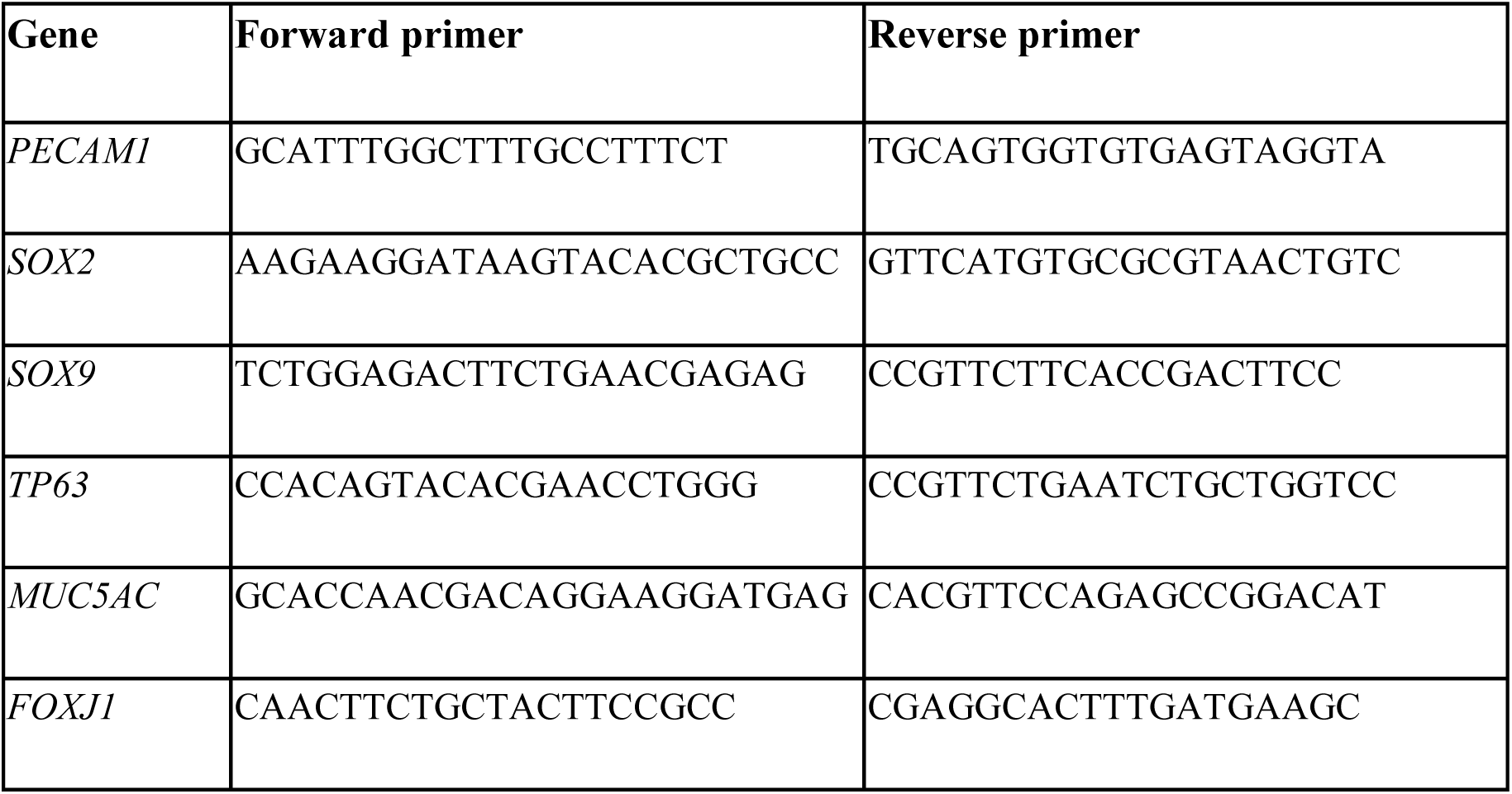
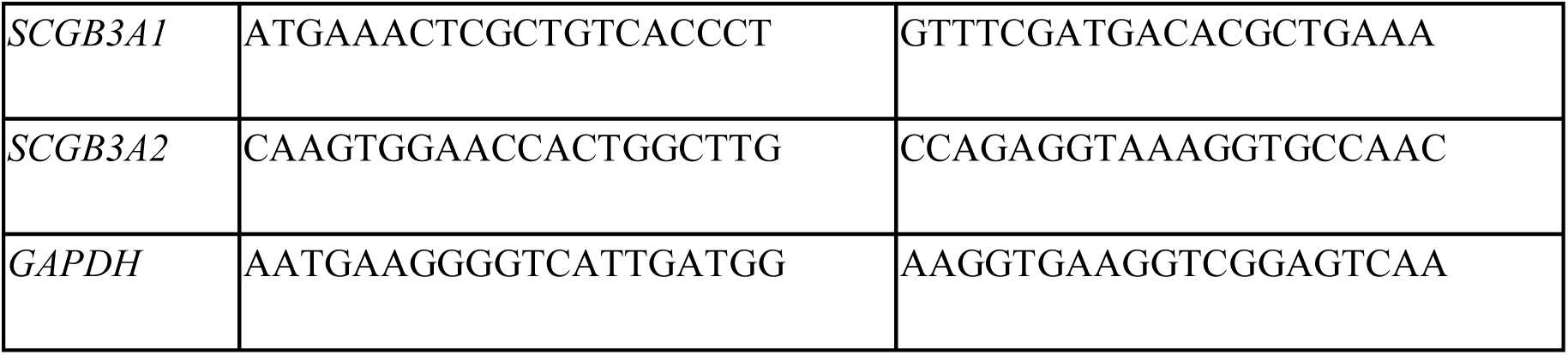
Oligonucleotide primers used for qRT-PCR

### Immunohistochemistry (cryosections)

Lung samples were fixed at 4°C in 4% paraformaldehyde (PFA) overnight. Samples were cut (if needed) to fit 15×15×5 mm molds. Post-fixation, samples were washed in PBS at 4°C, then sucrose-protected by incubation in 15%, 20%, then 30% sucrose in PBS for 1 hr each at room temperature. Samples were incubated 1:1 in 30% sucrose: optimal cutting temperature compound (OCT, Sakura) overnight at 4°C (small tissue fragments had an additional 100% OCT wash for 2 hrs at room temperature), then embedded in OCT and frozen on dry ice in an isopentane bath. 7 µm cryosections were cut, mounted onto SuperFrost® slides (VWR®) and stored at −80°C.

Sections were dried at room temperature for 2 hrs prior to staining. Tissue was permeabilized using 0.3% Triton-X100 in PBS for 10 min at room temperature, and slides were then washed in PBS. Where required, antigen retrieval was performed, by heating slides in 10 mM sodium citrate buffer pH6 in a full power microwave for 5 min, followed by 1 hr cooling at room temperature. Sections were blocked in 5% serum (same species as that of the secondary antibody) plus 1% BSA (bovine serum albumin), 0.1% Triton-X100 in PBS for 1 hr at room temperature. Primary antibodies (**Table S3**) were diluted in block solution and added to sections, with overnight incubation at 4°C. After PBS washes, sections were incubated in secondary antibodies (1:1000; all ThermoFisherScientific, **Table S3**) diluted in 5% serum (same as block), 0.1% Triton-X100 in PBS for 3 hrs at room temperature. Sections were incubated with DAPI (4′,6-diamidino-2-phenylindole, Merck) for 20 min, followed by PBS washes, then mounted in Fluoromount™ Aqueous Mounting Medium (Merck). Images were obtained with a Zeiss Axiophot or Leica DMi8 microscope.

### Immunohistochemistry (paraffin sections)

Lung samples were fixed at 4°C in 4% paraformaldehyde (PFA) overnight. Samples were cut (if needed) to fit molds. Post-fixation, samples were washed in PBS at 4°C, then incubated in 25% ethanol (EtOH) in 0.1% Tween-20 in PBS (PBS-T) for 30 min at 4°C. Further incubations in 50% EtOH/PBS-T, then 70% EtOH/PBS-T for 30 min each were performed, prior to transfer of the sample into an automated vacuum tissue processor for further dehydration and impregnation of the tissue with paraffin (Leica TP1050). Lungs were then embedded in paraffin wax. 3 µm sections were cut for immunohistochemistry and mounted onto SuperFrost® slides.

Sections were dewaxed using an automatic slide stainer (DRS 2000, Sakura) and then the same permeabilisation and staining procedure was used as for cryosections.

### Immunohistochemistry Quantification

A Zeiss Axioscan Z1 slide scanner was used to scan immunohistochemically stained tissue cryosections. *ImageJ* software was used to quantify the proportion of positively stained cells per DAPI-positive nucleus within each section. The mean ± SEM (standard error of mean) of three tissue sections was calculated for each sample and a minimum of three biological replicates were analyzed at each time point. p-values were calculated by one-way ANOVA followed by Tukey’s post-hoc test.

### smFISH/RNAscope

Fetal lung tissue was prepared as FFPE or fixed frozen blocks. For FFPE, samples were fixed in 10% neutral buffered formalin and embedded in wax as per standard protocols. For fixed frozen, samples were fixed overnight in cold 4% PFA, sucrose-protected as previously described, and frozen in OCT. To determine the best samples for analysis, morphology was assessed using hematoxylin and eosin staining and RNAscope 3-plex positive and negative control probes run. For RNAscope, 5-(FFPE) or 10-(fixed frozen) µm thick sections were cut on to SuperFrost® Plus slides and processed using the RNAscope 2.5 LS multiplex fluorescent assay (ACD, Bio-Techne) on the Leica BOND RX system (Leica). FFPE slides were baked and dewaxed (BOND protocol), incubated with protease III for 15 min, ER2 for 15 minutes at 95°C, whereas fixed frozen slides were first fixed for 15min with 4°C 4% PFA, baked for 30 min at 60°C, dehydrated through a standard ethanol series and run on the Leica BOND RX with 15 min protease III at room temperature and epitope retrieval for 5 min with ER2. All slides received opal 520, 570 and 650 fluorophores (Akoya Biosciences) at 1:1000 concentration. Probes used can be found in **Table S4** (ACD, Bio-Techne). Slides were counterstained with DAPI (ThermoFisher D1306, 300nM working dilution), and imaged on a Leica SP8 confocal or Perkin Elmer Opera Phenix. For some probe mixes, EpCAM antibody staining (abcam ab71916, 1:1000) was run following RNAScope, using TSA-biotin (1:400) and goat anti-rabbit IgG - HRP (1:1000).

### Digestion of human lung tissue and Flow Cytometry or FACS

The protocol for digesting lung tissue was optimized in our lab, based on methods originally used for digesting skin^64,65^. Lung tissue was dissected where necessary, to remove the trachea/any non-lung tissue, and weighed. The tissue was minced finely with scissors to create a paste and then resuspended in 3 to 8 ml (dependent on mass of tissue) digestion cocktail (composed of 2 mg/ml collagenase from *Clostridium histolyticum* (C9407), 0.5 mg/ml hyaluronidase (H3506) and 0.1 mg/ml DNase I (DN25)(all Merck) in AD+ medium: Advanced DMEM/F12 (12634010) with 10 mM HEPES (15630-056), 1X GlutaMax (35050038), 100 U/ml Penicillin and 100 μg/ml Streptomycin (all ThermoFisher Scientific)). Tissue was incubated at 37°C for 45 min, shaking at 400 rpm, then neutralized with AD+ medium and shaken vigorously for 30 sec. The sample was filtered through a 100 µM cell strainer, followed by a 40 µM cell strainer. For samples aged 11 pcw or older, red blood cell (RBC) depletion was performed, using EasySep™ RBC Depletion Reagent (StemCell Technologies, 18170), according to the manufacturer’s instructions. Cell number and viability were determined using Trypan Blue. Cells were blocked, to prevent non-specific binding, using Human TruStain FcX (Biolegend) and then stained with FACS antibodies (**Table S5**), according to the manufacturer’s instructions. Either Zombie UV™ fixable viability dye (Biolegend) or DAPI was utilized to stain and identify viable cells. Appropriate ‘Fluorescence Minus One’ (FMO) controls were included in analysis, to allow effective identification and gating of populations. Samples were either used for flow analysis or FACS. Flow data were collected using a BD LSRII and analyzed using Flowjo v10. FACS was performed using a BD FACS Aria.

FACS was utilized to isolate CD45^+^ immune cells, macrophages or dendritic cells, for downstream applications.

### Single-cell RNA sequencing (scRNAseq) and CITE-seq staining for single-cell proteogenomics

For scRNAseq, FACS-sorted CD45^+^ fetal lung cells were collected at the following time points and frozen, due to logistics: 8-9, 12 and 20 pcw. Cell suspensions were thawed rapidly at 37°C in a water bath. Warm RPMI1640 medium (ThermoFisher Scientific) (20 to 30 ml) containing 10% FBS (RPMI-FBS) was added slowly to the cells before centrifuging at 300 g for 5 min. This was followed by a wash in 5 ml RPMI-FBS. Cell number and viability were determined using Trypan Blue. Cells from different donors were pooled in equal numbers where possible. At this point, samples were either loaded directly onto a 10x chip, as described below, or stained for CITE-seq.

For CITE-seq staining, the reagent volumes were adjusted according to pooled cell number. For 5x10^5^ cells, samples were resuspended in 25 µl cell staining buffer (BioLegend, 420201). 2.5 µl Human TruStain FcX block (BioLegend, 422301) was added, and cells were incubated on ice for 10 min, to block non-specific binding. The cells were then stained with TotalSeq-C antibodies (BioLegend, 99814; full antibody list available in Stephenson *et al.*^66^) according to the manufacturer’s instructions. After incubating with 0.5 vials of TotalSeq-C for 30 min at 4 °C, cells were washed three times by centrifugation at 500 g for 5 min at 4 °C. Cells were counted again and processed immediately for 10x 5′ single cell capture (Chromium Next GEM Single Cell V(D)J Reagent Kit v1.1 with Feature Barcoding technology for cell Surface Protein-Rev D protocol). One lane of 4,000 to 25,000 cells were loaded per pool onto a 10x chip.

### Single-cell library construction and sequencing

Single-cell gene expression libraries (GEX) and V(D)J libraries were built using the 10X Chromium Single Cell V(D)J Kits (v1). The manufacturer’s protocol (10X Genomics) was followed using individual Chromium i7 Samples Indices. γδ TCR V(D)J libraries were prepared as previously described^67^. GEX and V(D)J libraries were pooled in 10:1 ratio and sequenced on a NovaSeq 600 S4 Flowcell aiming for a minimum of 50,000 paired-end (PE) reads per cell for GEX libraries and 5,000 PE reads per cell for V(D)J libraries. To maximize cell type and cell-state discovery, we combined these data, composed of 61,757 cells, with the immune compartment of our previous unbiased single-cell whole fetal lung scRNAseq data, which added an additional 15,802 cells^21^, to give a total of 77, 559 cells.

### DNA extraction and genotyping

Fetal skin samples and corresponding maternal salival samples were used for DNA extraction, according to the manufacturer’s protocol (Qiagen, DNeasy Blood and Tissue Kit, 69504, and QIAamp DNA Investigator Kit, 56504 respectively). DNA samples were used for genotyping, performed with the Affymetrix UK Biobank Axiom™ Array kit by Cambridge Genomic Services (CGS).

### Single-cell data quantification

scRNAseq data were mapped with STARsolo 2.7.3a^68^ using the GRCh38 reference distributed by 10X, version 3.0.0 (derived from Ensembl 93). Cell calling was performed with an implementation of EmptyDrops extracted from Cell Ranger 3.0.2 (distributed as emptydrops on PyPi). For single-cell V(D)J data, reads were mapped with Cell Ranger 4.0.0 to the 10X distributed VDJ reference, version 4.0.0. Single-cell CITE-seq data was mapped with Cell Ranger 4.0.0 to the 10X distributed GRCh38 reference, version 3.0.0.

### Genetic demultiplexing of donors

To identify the donor identities of each cell in multiplexed samples, souporcell^69^ (command for a 2-plex sample: singularity exec −B $PWDsouporcell/souporcell.sif souporcell_pipeline.py -i Aligned.sortedByCoord.out.bam -b barcodes.tsv -f refdata-cellranger-GRCh38-3.0.0/fasta/genome.fa - t 15 -o souporcell_known --known_genotypes jbID-hg38.vcf --known_genotypes_sample_names JB12 JB16 --skip_remap True -k 2) was used to match our genotyping results of each donor to empirical genotypes for each cell inferred from the scRNAseq reads. Cells with status “singlet” were each assigned a donor while others were labeled with mixed donors.

### Single-cell RNA-seq and CITE-seq data downstream analyses

After demultiplexing, gene expression together with CITE-seq antibody barcode count matrices from the Starsolo-EmptyDrops pipeline and filtered matrices from the Cell Ranger-SoupX pipeline described above were loaded and concatenated. Each row (cell) of the concatenated matrix was then divided by the total number of non-antibody non-mitochondria UMI counts (referred to as “total UMI counts” below) of the corresponding cell, and multiplied by a scaling factor of 10,000, followed by a natural log transformation (pseudocount = 1). Cells with fewer than 200 genes expressed or more than 20% of reads mapped to mitochondrial sequences were removed. Genes that were expressed in fewer than 5 cells were discarded.

The resulting matrix was then concatenated with the immune-cell subset of our previously generated single-cell data. For each sample, highly variable genes were calculated using the default setting of Scanpy 1.5.0. Among these, highly correlated genes were selected for each sample as described in He et al. 2022^21^, and those selected in at least three samples were used as feature genes. The feature gene count matrix was regressed out against cell cycle scores (S and G2M scores calculated according to Scanpy’s instruction), total UMI counts and fraction of mitochondria reads. The residue matrix was scaled along the gene dimension and used for principal component analysis (PCA). To mitigate technical effect due to different cell-isolation protocols used for our previously and newly generated datasets (indicated by a boolean variable ‘project’), BBKNN was used to integrate the datasets together (batch_key=’project’,n_pcs=50, neighbors_within_batch=10). The resulting nearest neighbor graph was used for Leiden clustering and UMAP embedding.

Clusters were annotated based on marker genes and literature. The clusters with cells from only one sample or non-informative/house-keeping genes as marker genes were flagged as technical clusters and discarded. The aforementioned steps from regression to discarding artifact cell clusters were repeated once more to remove residue technical clusters.

Background and non-specific staining by the antibodies used in CITE-seq was estimated using SoupX (v.1.4.8), which models the background signal on near-empty droplets. The soupQuantile and tfidfMin parameters were set to 0.25 and 0.2, respectively, and lowered by decrements of 0.05 until the contamination fraction was calculated using the autoEstCont function. Antibody-derived tag counts were normalized with the centered log-ratio transformation.

### Maternal cell inference

Maternally-derived cells were inferred using two methods. The libraries containing one single donor were selected and fed into the souporcell 2-plex sample pipeline without known genotypes specified, in order to cluster cells into fetal and maternal cells based on genetic background. Additionally, we picked male samples (high expression of DDX3Y) and evaluated the expression levels of XIST cell type by cell type. Neither method suggested evidence of prominent maternal cell presence.

### Differential gene expression analysis

To characterize maturation of T cells through fetal and early pediatric life, we tested for differential gene expression in naive T cells from healthy pediatric donors from Yoshida *et al*^35^. Differential expression across age within these naive CD4 or CD8 T cells was tested by fitting a gamma poisson generalized linear model on log_2_ transformed age and by creating pseudo-bulks of each donor with the glmGamPoi package. Fetal and pediatric expression of the top 25 most significant upregulated and downregulated genes are shown in **Fig S5F**.

### Cell type composition analysis

The number of cells for each sample and cell type composition was modeled with a generalized linear mixed model with a Poisson outcome using the ‘glmer’ function in the lme4 R package. The donor ID, sequencing batch, and fetal age were fitted as random effects to overcome the colinearity among the factors. The effect of each factor on cell type composition was estimated by the interaction term with the cell type. The standard error of variance parameter for each factor was estimated using the numDeriv package. The conditional distribution of the fold change estimate of a level of each factor was obtained by the ‘ranef’ function in the lme4 package. The statistical significance of the fold change estimate was measured by the local true sign rate (LTSR), which is the probability that the estimated direction of the effect is true, that is, the probability that the true log-transformed fold change is greater than 0 if the estimated mean is positive (or less than 0 if the estimated mean is negative). It is calculated on the basis of the estimated mean and s.d. of the distribution of the effect (log-transformed fold change), which is to an extent similar to performing a (one-sided) one-sample *Z*-test and showing (1 − *P*).

### B cell trajectory analyses

The soup-corrected single cell transcriptomics data was preprocessed using Scanpy^70^ version 1.8.1. In particular, the cell cycle effect and fraction of mitochondrial reads were regressed out using sc.pp.regress_out() and batch correction was performed using BBKNN^26^. PAGA^71^ was applied with sc.tl.paga() to examine the connectivities between cell types, before the final UMAPs were computed using the results of PAGA for initialisation as described in^71^. Data and UMAPs were exported to R for running monocle3^72,73^ to find a principal graph and define pseudotime. Differentially expressed genes along pseudotime were then computed using a graph-based test (morans’ I)^73,74^ over the principal graph. This allows us to identify genes that are upregulated at any point in pseudotime. The results were visualized using the complexHeatmap^75^ and seriation^76^ packages in R, after smoothing gene expression with smoothing splines (smooth.spline(), df=12).

### BCR and TCR analyses

Single-cell BCR and TCR data were initially processed with cellranger-vdj (v.6.0.0). Contigs contained in all_contigs.fasta and all_contig_annotations.csv were then processed further using *dandelion*^66^ singularity container (v.0.2.1) (https://www.github.com/zktuong/dandelion). Pairwise wilcoxon rank sum test was performed on BCR heavy/light chain CDR3 junction lengths (nucleotide) using *scipy.stats.ranksums* (v1.5.2). Benjamini-Hochberg false discovery rate correction was applied using *statsmodels.stats.multitest.fdrcorrection* (v0.12.1). Significance testing was performed using mean values of each sample between cell types at each time point.

Single-cell γδTCR data was initially processed with cellranger-vdj, using the 10X-distributed GRCh38 VDJ reference (both v.4.0.0). High confidence contigs contained in all_contigs.fasta and all_contig_annotations.csv were then processed further using dandelion^66^.

In nonproductive TCR analysis, all_contig_annotations.csv from cellranger-vdj was used as input for αβTCR; while all_contig_igblast_db-all.tsv from dandelion was used for gdTCR. Only high confidence contigs were used. For each unique cell barcode, the primary V(D)J contig was determined based on its ‘productive’ status (’productive’ contig would take priority over ‘nonproductive’ contig) followed by their umi count. The majority of ‘nonproductive’ contigs identified were contigs with J and C gene segments, without any V segment.

### Integrated analysis with cross-organ immune atlas

The cross-organ dataset of blood and immune cells from Suo et al. 2022 was downloaded from https://developmental.cellatlas.io/fetal-immune. We concatenated gene expression profiles from the 9 tissues and lung and integrated the datasets using scVI^77^ using the implementation in the python package scvi-tools 0.14.5 (model parameters: n_latent=20, encode_covariates=True, dropout_rate=0.2, n_layers=2; training parameters: early_stopping=True, train_size=0.9, early_stopping_patience=45, max_epochs=400, batch_size=1024, limit_train_batches=20). For the differential cell abundance analysis on neighborhoods we used the Milo framework^78^, as implemented in the python package milopy v0.1.0. For this analysis, we excluded lung samples older than 17 pcw, to match the age range in the cross-tissue dataset, and tested for significant increase or decrease in cell numbers in lung samples compared to all other tissues. The effect of FACS sorting protocol on cell abundance was accounted for in the differential abundance model. For differential expression analysis on lung-enriched ILC and NK neighborhoods, we label cells belonging to ILC or NK lung-enriched neighborhoods and compare them to cells belonging to other ILC and NK neighborhoods. We performed differential expression analysis pseudo-bulking by sample using the R package glmGamPoi. When selecting DE genes, we excluded genes that were found to be differentially expressed between lung-enriched and other neighborhoods, a set of control cell types where we don’t expect to see lung-specific signatures (T progenitors, HSC/MPP, Immature B cells, Treg).

### Microdissection of epithelial tip cells from embryonic lung

Human embryonic lung epithelial tip organoids were established and maintained as previously described^6^. Briefly, embryonic fetal lung lobes were incubated in Advanced DMEM/F12 medium supplemented with 8 U/ml Dispase (ThermoFisher Scientific, Gibco) for 2 min at room temperature. Mesenchyme was carefully dissected away using tungsten needles. Epithelial tips were micro-dissected by cutting the very end of branching epithelial tubes under a dissection microscope. Tips were transferred into 25 μl of Matrigel (Corning, 356231) and plated in a 48 well low-attachment plate (Greiner). The plate was incubated for 5 min at 37°C to solidify the Matrigel before adding 250 μl of self-renewing medium (**Table S6**) per well. The plate was incubated under standard tissue culture conditions (37°C, 5% CO2).

### Maintenance of 3D human embryonic lung organoids

Organoids were cultured in Matrigel (Corning, 356231) with self-renewing medium. Medium was changed twice a week, and organoids were passaged every 10-14 days. Organoid lines between passage 3 and passage 7 were used for the following functional experiments.

### Cytokine treatment of organoids

10 ng/ml Recombinant Human IL-1α (Peprotech, 200-01A), IL-1β (Peprotech, 200-01B), IL-4 (Peprotech, 200-04), IL-6 (200-06), IL-13 (Peprotech, 200-13), IL-17A (Peprotech, 200-17), IL-22 (Peprotech, 200-22), IFN-γ (Peprotech, 300-02), TNF-α (Peprotech, 300-01A), TNF-β (Peprotech, 300-01B) were added to the self-renewing medium for 3, 7 and 14 days. Cytokine/self-renewing medium was changed twice-weekly. To inhibit IL-1 signaling, 100 ng/ml of IL-1Ra (Peprotech, 200-01RA) or 2 μM (5Z)-7-Oxozeaenol (Bio-techne, 3604) was added to the self-renewing medium.

### Dual SMAD activation medium

Self-renewing medium without Noggin and SB431542 was supplemented with 100 ng/ml TGF-β1 (Peprotech, 100-21) and 100 ng/ml BMP4 (R&D Systems, 314-BP). This was termed Dual SMAD activation medium^44^. To promote TP63^+^ basal cell differentiation, self-renewing medium was removed and changed to Dual SMAD activation medium for 3 days.

### Recovering organoids from Matrigel for RNA extraction, protein extraction or immunostaining

Organoids were removed from Matrigel using Matrigel Cell Recovery Solution (Corning, 354253). After removing the culture medium from the wells and washing with PBS, 1 ml of ice-cold Matrigel Cell Recovery Solution was added to each well and incubated for 45 min at 4°C. Organoids were washed with ice-cold PBS and spun down at 200 g at 4°C.

### Quantitative PCR of organoids

Organoids removed from Matrigel were lysed using 350 μl of RLT buffer. Total RNA was extracted using the RNeasy Mini Kit (Qiagen) according to manufacturer’s instructions. RNA yields were measured using Nanodrop (ThermoFisher Scientific). 0.5 μg RNA was reverse transcribed using qScript cDNA Supermix (Quantabio, 733-1177), following the manufacturer’s protocol. cDNA was used for qPCR reactions using Power SYBR Green Master Mix (Life Technologies Ltd). Relative gene expression was calculated using the ΔΔCT method relative to GAPDH control. P-values were obtained using an unpaired two-tailed student’s t-test with unequal variance. For primer sequences see **Table 1**.

### Western blot

Organoids removed from Matrigel were lysed in ice-cold RIPA buffer (ThermoFisher Scientific, 89900) containing protease inhibitor cocktail (Roche, 589297001) and phosphatase inhibitors (Roche, 4906845001). Total protein concentration was quantified by means of BCA protein assay (Merck). 10-20 μg of protein was separated by size using a 4-12% Bis-Tris Plus gel ThermoFisher Scientific) and transferred to a nitrocellulose membrane, using an iBlot2™ transfer stack (ThermoFisher Scientific, IB23002) with the iBlot2™ Gel Transfer Device (ThermoFisher Scientific). The membrane was incubated with specific primary antibody overnight at 4°C, HRP-linked secondary antibody (Cell Signaling Technologies) for 1 hour at room temperature, followed by chemiluminescence detection (Immobilon® Crescendo HRP Substrate, Millipore) with the ImageQuant LAS 4000 (GE Healthcare).

### Whole mount staining of organoids

Organoids removed from Matrigel were fixed with 4% paraformaldehyde (PFA) for 30 min at 4°C. After washing in PBS, organoids were transferred to a 48 well plate for staining. Permeabilization was performed in 0.5% (v/v) Triton-X100 in PBS for 30 min at room temperature, followed by blocking in 1% BSA, 5% normal donkey serum, 0.2% Triton-X100 in PBS (blocking solution) for 1 hr at room temperature. Organoids were incubated with primary antibodies in blocking solution overnight at 4°C. After washing off primary antibodies, secondary antibodies (1:1000 dilution) in blocking solution were added and incubated overnight at 4°C. After washing, DAPI (Thermo scientific, 62248) staining was performed for 30 min at 4°C, followed by clearing/mounting using 2’-2’-thio-diethanol (Sigma, 166782) as previously described^6^. Confocal images were acquired using an SP8vis microscope (Leica).

### Organoid colony forming assay

Organoids removed from Matrigel were transferred to 15 ml tubes. Organoids were resuspended in 1 ml TrypLEexpress (Gibco, 1265036) and incubated at 37°C for up to 10 mins, triturating gently every 2 mins with a P1000 pipette. 9 ml of 10% FBS was added, followed by centrifugation at 4°C (300 g, 5 min). The pellet was resuspended in 1% BSA in HBSS and filtered through 40μm cell strainers into 50 ml tubes. The single cell suspension was centrifuged as before and resuspended in HBSS. Cells were counted using Trypan Blue, to check single cell efficiency and viability. Cells were seeded into a 96-well plate (500 cells/well)(Greiner) in a 50 μl Matrigel droplet. 100 μl of self-renewing medium supplemented with 10 nM Rho kinase inhibitor Y-27632 (Merck, 688000) was added for 2 days, and then changed to SR medium supplemented with either 10 ng/ml IL-1β or 100 ng/ml IL-1Ra and cultured for a further 7 days before counting and calculating organoids size and organoid forming efficiency using *ImageJ* software.

### Macrophage culture

Macrophages (HLA-DR^+^CD206^+^CD169^−^) and monocytes (HLA-DR^+^CD206^−^CD169^−^) were isolated from human fetal lung tissue by FACS (**Fig S1C**). Isolated macrophages and monocytes were cultured in TexMACS medium (Miltenyi Biotec, 130-097-196) supplemented with 100 ng/ml human recombinant M-CSF (Peprotech, 300-25) for 7 days. On day 3, 5 and 7 of the culture, supernatant was removed/stored and replaced with fresh medium.

### Dendritic cell culture

Dendritic cells were isolated from human fetal lung tissue by FACS (**Fig S1C**). Isolated dendritic cells were cultured in ImmunoCult-ACF DC Medium (STEMCELL Technologies, 10986) supplemented with ImmunoCult DC Maturation Supplement (STEMCELL Technologies, 10989) for up to 7 days. On day 3, 5 and 7 of the culture, supernatant was removed/stored and replaced with fresh medium.

### Cytokine array

Supernatants were harvested from cultured immune cells and centrifuged at 1,000 g to remove debris. Cytokines secreted into the supernatant were analyzed using a Human Cytokine Antibody Array (abcam, ab133997), according to the manufacturer’s instructions.

## Acknowledgements

This publication is part of the Human Cell Atlas - www.humancellatlas.org/publications. We are grateful to the Joint MRC/Wellcome Trust (grant # MR/R006237/1) Human Developmental Biology Resource (www.hdbr.org) for provision of human embryonic and fetal material. We acknowledge Jamie Evans for his assistance at the UCL Rayne flow cytometry core facility. We acknowledge the Cellular Genetics IT and Phenotyping group, New Pipeline Group, Spatial Genomics Platform (in particular Sophie Pritchard) and the DNA pipelines of Wellcome Sanger Institute. We are grateful to Cambridge Genomic Services, Department of Pathology, University of Cambridge for carrying out genotyping work. We also acknowledge assistance from Stuart Forbes, Davide Danovi and Niwa Ali. M.Z.N. acknowledges funding from a MRC Clinician Scientist Fellowship (MR/W00111X/1), Rosetrees Trust (M899) and the Longfonds BREATH consortium. M.Z.N. and J.L.B. acknowledge funding from the Rutherford Fund Fellowship allocated by the MRC UK Regenerative Medicine Platform 2 (MR/5005579/1). M.Y. was funded by The Jikei University School of Medicine and Action Medical Research (GN2911). K.B.W. acknowledges funding from University College London, Birkbeck MRC Doctoral Training Programme. K.B.M. and S.A.T. are supported by Wellcome (WT211276/Z/18/Z and Sanger core grant WT206194). K.B.M. and E.L.R. acknowledge funding from the MRC (MR/S035907/1). E.L.R. acknowledges funding from the MRC (MR/P009581/1). Z.K.T. and M.R.C. are supported by a Medical Research Council Research Project Grant (MR/S035842/1). M.R.C. is supported by a National Institute of Health Research (NIHR) Research Professorship (RP-2017-08-ST2-002), a Wellcome Investigator Award (220268/Z/20/Z), the Blood and Transplant Research Unit in Organ Donation and the NIHR Cambridge Biomedical Research Centre.

## Author Contributions

M.Z.N., K.B.M. and S.A.T. conceived, set up and directed the study. J.L.B., M.Y. and K.B.W. performed 10X and CITE-seq. J.L.B. performed tissue digestion, cell sorting, DNA extraction for genotyping and immunohistochemistry. P.H., R.G.H.L., C.S., E.D., J.P.P., A.O., A.P., Z.K.T., K.P., M.Y. and K.B.W. performed bioinformatic analysis. P.H. annotated cell types. M.Y., K.B.W., I.T.H. and J.A.H. performed functional organoid experiments. M.Y. performed the immune cell culture, analysis and immunohistochemistry. J.L.B., K.B.W. and J.A.H. performed qPCR. A.W.C. performed smFISH/RNAscope staining and imaging. L.M. and L.R. prepared sequencing libraries and conducted the sequencing. P.A.L. performed genotype calling. R.E.H. and V.H.T. provided support in student supervision (to K.B.W.). J.R. provided immunological expertise. S.M.J. provided regular intellectual input throughout the study. K.L., D.S. and E.L.R. provided additional whole fetal lung single cell data. M.H. provided expertise on the development of the immune system. J.M. and S.A.T. provided supervision (to P.H.). M.C. provided supervision (to Z.K.T.). J.L.B., P.H., M.Y., C.S., Z.K.T., R.G.H.L., K.B.W., K.B.M and M.Z.N. wrote the manuscript. All authors read and/or edited the manuscript.

## Data Availability

Raw FASTQ and processed BED formatted data will be deposited on ArrayExpress and BioStudies under accession number E-MTAB-11528.

## Supplementary Table Titles

**Table S1: Marker genes** related to **Figure 2 and 3**.

**Table S2: Confusion matrix, fetal lung immune vs pan-fetal immune** related to **Figure S5**.

**Table S3:** Primary and secondary antibodies for IHC related to Methods.

| **Antibody** | **Host species** | **Company** | **Catalogue number** | **Final dilution** | **Antigen retrieval?** |
| --- | --- | --- | --- | --- | --- |
| **CD45** | rabbit | abcam | ab10558 | 1:200 |  |
| **CD45** | mouse | Tonbo Biosciences | 70-0459 | 1:500 |  |
| **CD31** | mouse | abcam | ab9498 | 1:200 |  |
| **CD31** | rabbit | GeneTex | GTX81432 | 1:50 | Y |
| **ECAD** | mouse | BD Transduction Laboratories™ | 610182 | 1:500 |  |
| **ECAD** | rat | ThermoFisher Scientific | 13-1900 | 1:500 |  |
| **ECAD** | goat | R&D Systems™ | AF648SP | 1:20 |  |
| **CD4** | rabbit | abcam | ab133616 | 1:100 | Y |
| **CD8** | rat | Bio-Rad | MCA351G | 1:100 |  |
| **CD56** | rabbit | antibodies-online.com | ABIN1859967 | 1:50 |  |
| **CD20** | mouse | abcam | ab9475 | 1:100 | Y |
| **CD11c** | rabbit | abcam | ab53632 | 1:100 |  |
| **CD68** | mouse | GeneTex | GTX75390 | 1:100 |  |
| **SOX2** | mouse | abcam | ab79351 | 1:500 |  |
| **SOX2** | rat | ThermoFisher Scientific | 14-9811-80 | 1:500 |  |
| **SOX9** | rabbit | abcam | ab185966 | 1:500 |  |
| **p63** | rabbit | Cell Signaling Technology | 13109 | 1:200 |  |
| **IL-1R1** | mouse | Santa Cruz | sc-393998 | 1:50 |  |
| **IL-1RAcP** | mouse | Santa Cruz | sc-376872 | 1:50 |  |
| **CD206** | mouse | ThermoFisher Scientific | MA5-28581 | 1:50 |  |
| **CD1C** | mouse | abnova | MAB22942 | 1:200 |  |
| **Keratin 5** | chicken | Biolegend | 905901 | 1:500 |  |
| **Antibody** |  |  | **Catalogue number** |  |  |
| **Goat α-mouse Alexa Fluor 488** |  |  | A11029 |  |  |
| **Goat α-rabbit Alexa Fluor 488** |  |  | A11034 |  |  |
| **Goat α-mouse Alexa Fluor 555** |  |  | A21424 |  |  |
| **Goat α-rat Alexa Fluor 555** |  |  | A21434 |  |  |
| **Goat α-mouse Alexa Fluor 647** |  |  | A21236 |  |  |
| **Donkey α-rabbit Alexa Fluor 488** |  |  | A21206 |  |  |
| **Donkey α-mouse Alexa Fluor 488** |  |  | A21202 |  |  |
| **Donkey α-goat Alexa Fluor 647** |  |  | A21447 |  |  |
| **Donkey α-rabbit Alexa Fluor 555** |  |  | A31572 |  |  |
| **Chicken α-rat Alexa Fluor 647** |  |  | A21472 |  |  |

**Table S4:** smFISH probes related to Methods.

| **Probe** | **Catalogue number** |
| --- | --- |
| **AZU1** | 1130958-C2 |
| **BEST3** | 487188-C4 |
| **C19Orf77** | 407078-C2 |
| **CCDC81** | 1084658-C3 |
| **CCL19** | 47468 |
| **CCL21** | 474378-C4 |
| **CCL22** | 468708-C4 |
| **CCR10** | 313488-C4 |
| **CD1C** | 514768-C3 |
| **CLECL1** | 1044408-C2 |
| **CXCR5** | 4741778 |
| **DDX3Y** | 892688 |
| **DNTT** | 561978-C3 |
| **ELANE** | 509728-C2 |
| **FCER1A** | 311298-C3 |
| **IL17A** | 310938-C3 |
| **IL1B** | 310368 |
| **ITGA2B** | 435078-C2 |
| **MPO** | 603098-C3 |
| **MS4A1** | 426778-C2 |
| **NCR2** | 826958 |
| **PF4** | 531748-C3 |
| **PTPRC** | 470908-C2 |
| **PTPRC** | 601998-C3 |
| **PTPRC** | 601998-C4 |
| **RAG1** | 545758-C2 |
| **RORC** | 556998-C2 |
| **S100A12** | 820168-C3 |
| **S100A8** | 425278 |
| **S100A9** | 417588-C2 |
| **SCN1B** | 875838-C4 |
| **SPINK2** | 521688 |
| **TESPA1** | 865178 |
| **VPREB1** | 580608 |
| **VPREB3** | 1129068-C1 |
| **XIST** | 311238-C2 |

**Table S5:** FACS antibodies related to Methods.

| **Antibody** | **Conjugate** | **Company** | **Catalogue number** | **Final dilution** |
| --- | --- | --- | --- | --- |
| **CD45** | PE | BD Biosciences | 555483 | 1:20 |
| **CD45** | BV711 | Biolegend | 304049 | 1:50 |
| **CD31** | BV421 | Biolegend | 303124 | 1:50 |
| **CD31** | FITC | Biolegend | 303104 | 1:50 |
| **EpCAM** | APC | Biolegend | 324208 | 1:50 |
| **EpCAM** | FITC | Biolegend | 324203 | 1:50 |
| **CD235a** | FITC | Biolegend | 349104 | 1:50 |
| **CD4** | BUV395 | BD Biosciences | 344711 | 1:50 |
| **CD8** | PE-Cy7 | Biolegend | 344711 | 1:50 |
| **CD3** | APC-Cy7 | Biolegend | 300317 | 1:50 |
| **CD25** | BV421 | Biolegend | 302629 | 1:50 |
| **FOXP3** | PE | Biolegend | 320107 | 1:50 |
| **CD19** | BUV395 | BD Biosciences | 563549 | 1:50 |
| **CD56** | PE | Biolegend | 318305 | 1:50 |
| **HLA-DR** | APC-Cy7 | Biolegend | 307617 | 1:50 |
| **CD14** | PE-Cy7 | Biolegend | 301813 | 1:50 |
| **CD206** | BV421 | Biolegend | 321125 | 1:50 |
| **CD169** | APC | Biolegend | 346007 | 1:50 |
| **CD11c** | BUV395 | BD Biosciences | 563788 | 1:50 |
| **CD123** | PE | Biolegend | 306005 | 1:50 |

**Table S6:**
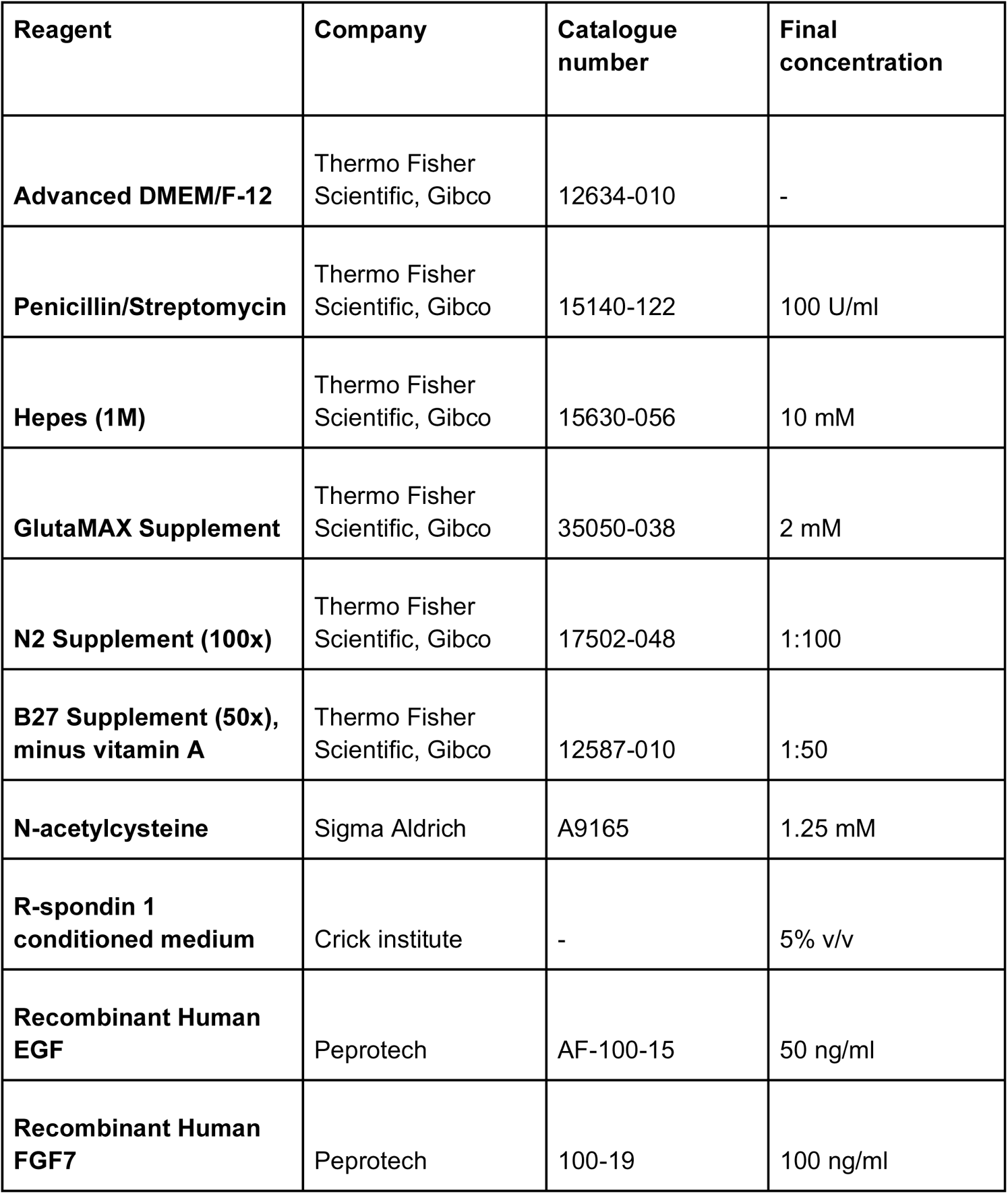
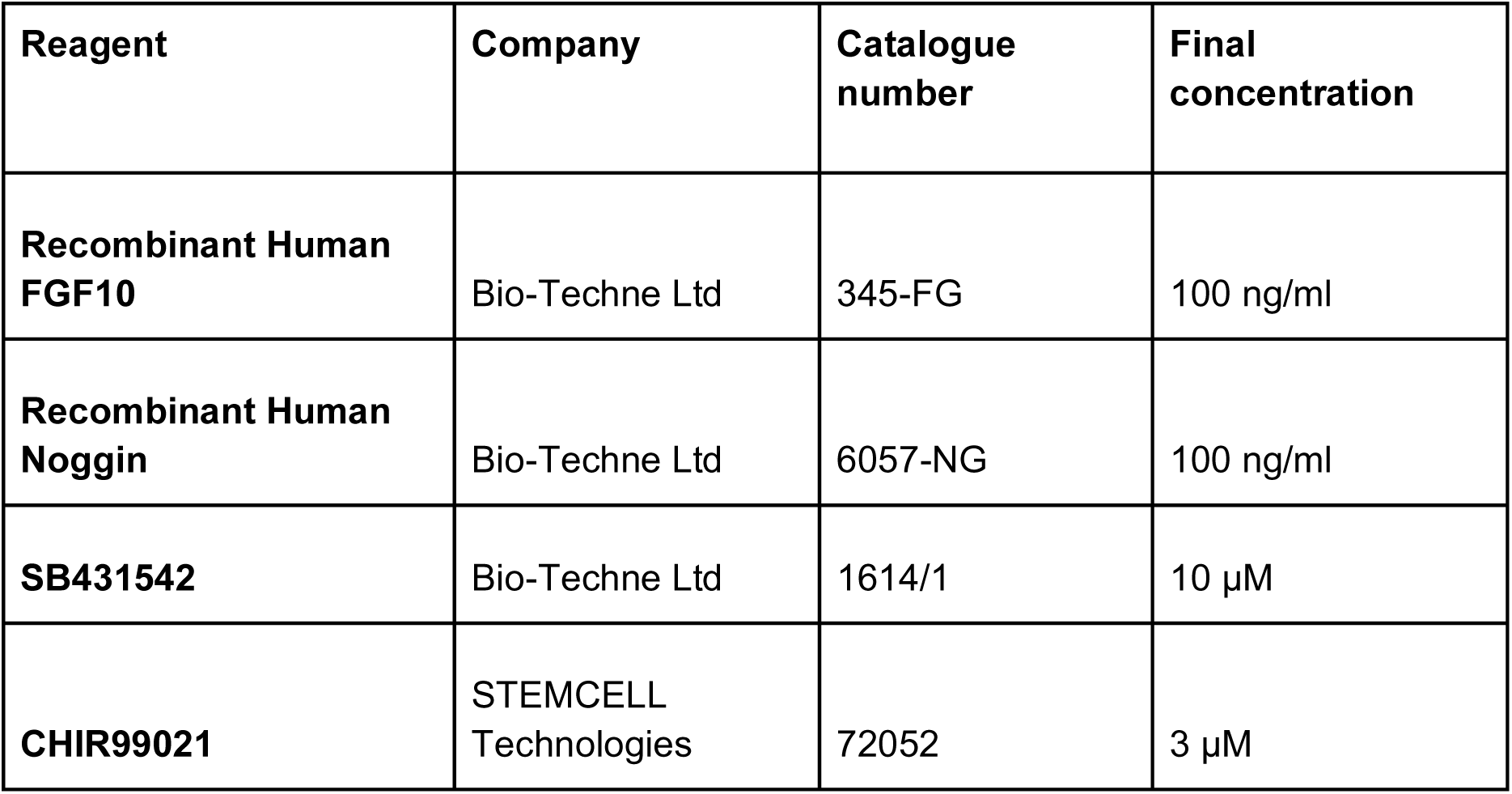
Self-renewing medium related to Methods.

## Supplementary Figure Legends

**Figure S1.**
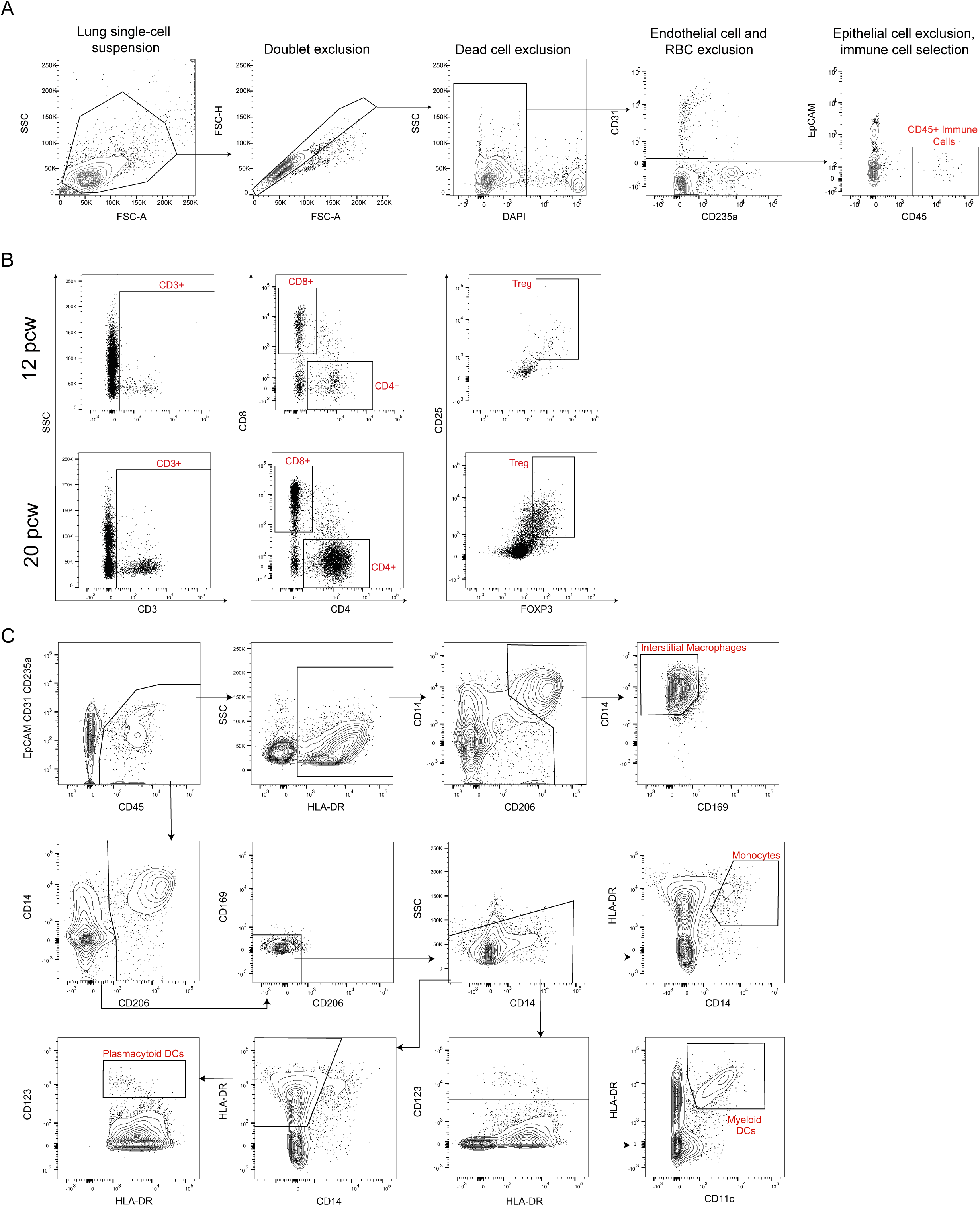
FACS gating strategies. Whole fetal lungs were digested, stained using specific antibody panels and immune cells were isolated by FACS or analyzed using flow cytometry. (**A**) shows the FACS gating strategy for isolating CD45^+^ immune cells in a representative 12 pcw sample. This strategy was used as the basis for all FACS and flow cytometry performed. Debris and cell aggregates were gated out of the single-cell suspension on the first plot, which shows FSC-A (forward scatter-area) versus SSC (side scatter). Doublets were excluded (FSC-A versus FSC-H (forward scatter-height)). From the singlets, DAPI^+^ or Zombie UV™^+^ dead cells were excluded. Endothelial cells (CD31^+^), red blood cells (RBCs, CD235a^+^) and epithelial cells (EpCAM^+^) were removed and CD45^+^ immune cells were collected. For T cell and myeloid cell analysis we combined CD31, CD235a and EpCAM markers into a ‘dump’ channel, whereby they were labeled with the same FITC fluorochrome. (**B**) shows T cells separated into the following populations in representative 12 and 20 pcw fetal lung samples (one million live cells were analyzed for each): the whole T cell population (CD3^+^), CD4^+^ T helper cells, CD8^+^ cytotoxic T cells and Tregs (CD3^+^CD4^+^FOXP3^+^CD25^+^). (**C**) shows the gating strategy for isolating myeloid cells in a representative 20 pcw lung sample. We added the following markers to the initial antibody panel: CD14, HLA-DR, CD206, CD169 and CD123, which allowed separation of interstitial macrophages, monocytes, myeloid DCs and plasmacytoid DCs. We designed this antibody panel based on previous studies^79–82^. Related to **Fig 1, 2** and **6**.

**Figure S2.**
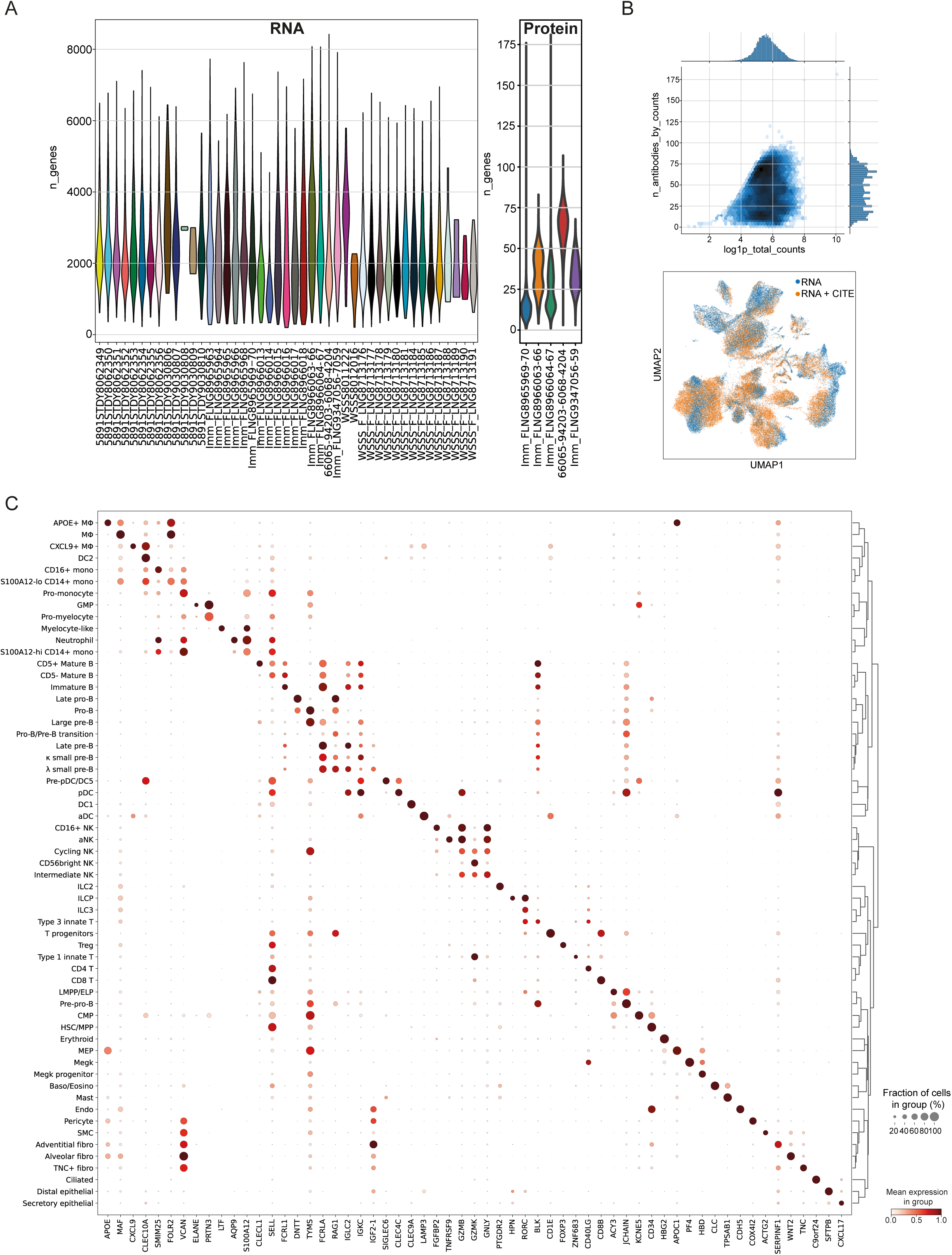
scRNAseq data quality control and marker gene expression. (**A**) A violin plot of the number of genes (left) or barcoded antibodies (right) detected per cell within each scRNAseq/CITE-seq library. (**B**) The number of antibodies detected plotted over the log-transformed total number of counts (upper) and the cells with or without CITE-seq measurement embedded on the transcriptomic UMAP (lower). Each dot represents one single cell. (**C**) The curated representative marker genes separating cell type/state clusters. Gene expression levels are scaled against the maximum for each gene. Related to Fig 2. See also **Table S1**.

**Figure S3.**
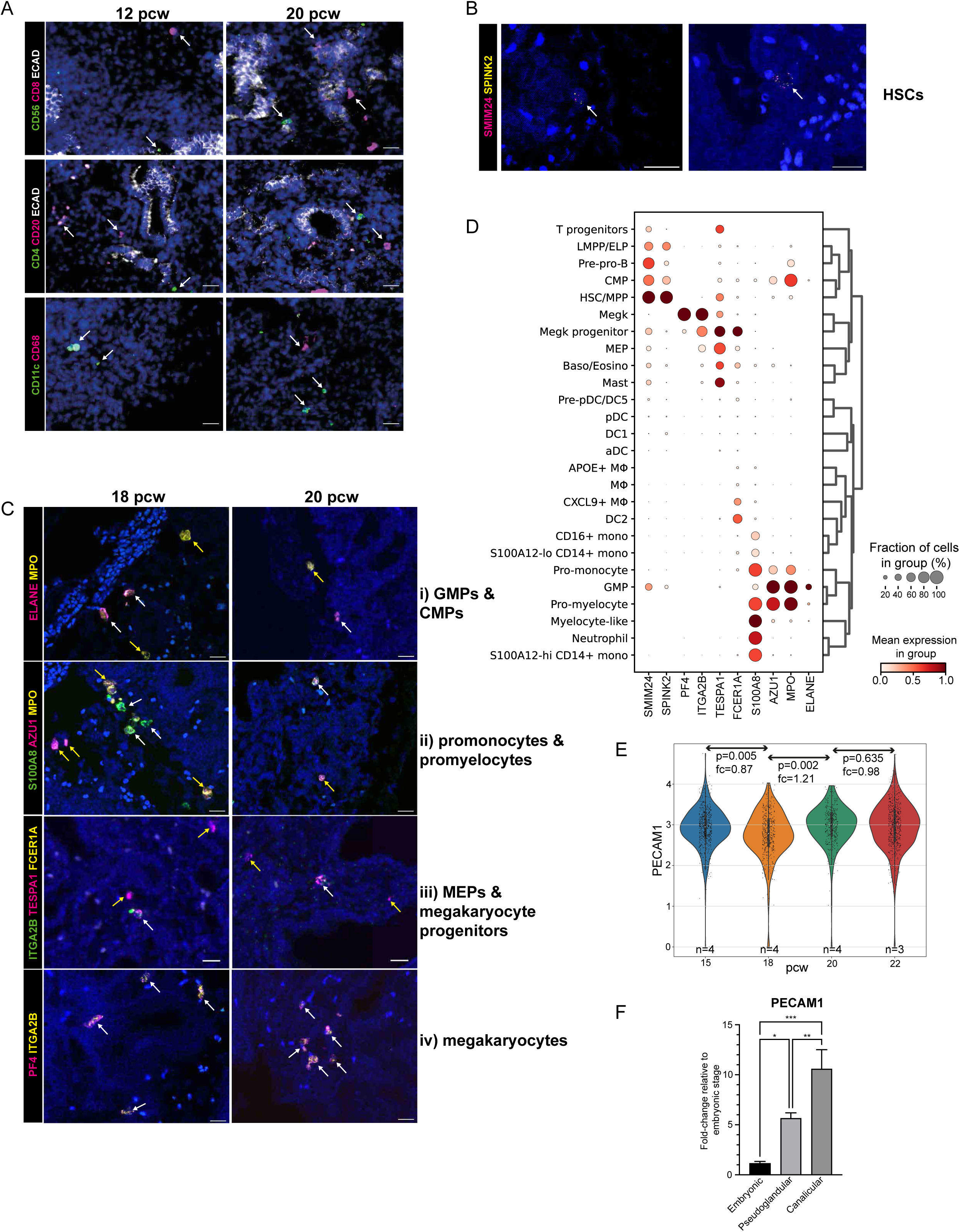
Validation of immune cell populations. (**A**) Representative IHC images show the presence of the following immune cell markers at 12 and 20 pcw: CD8, CD56, CD4, CD20, CD11c and CD68 (white arrows indicate cells expressing those markers). In the top two panels, the epithelium is stained with ECAD. (**B**) and (**C**) show representative RNAscope images of cell types identified in the scRNAseq dataset. (**B**) shows hematopoietic stem cells (HSCs, white arrows: SMIM24^+^SPINK^+^). (**C**) shows (**i**) white arrows: GMPs (ELANE^+^MPO^+^), yellow arrows: CMPs (MPO^+^), (**ii**) white arrows: promonocytes (S100A8^+^MPO^+^), yellow arrows: promyelocytes (AZU1^+^MPO^+^), (**iii**) white arrows: megakaryocyte progenitors (TESPA1^+^ITGA2B^+^FCER1A^+^), yellow arrows: MEPs (TESPA1^+^), (**iv**) white arrows: megakaryocytes (PF4^+^ITGA2B^+^). In all images, blue: DAPI^+^ nuclei; scale bar = 20µM. (**D**) shows cell type-specific expression of probed genes. (**E**) Distribution of log-transformed *PECAM1* (*CD31*) transcript counts per age group (values and fold changes are based on two-tailed t-tests of sample means). (**F**) shows expression of the endothelial marker *CD31* determined by realtime PCR, using RNA from whole fetal lungs at the embryonic (7-8 pcw), pseudoglandular (10-16 pcw) and canalicular (19-20 pcw) stages of development (n≥3, mean ± SEM). Related to Fig 2.

**Figure S4.**
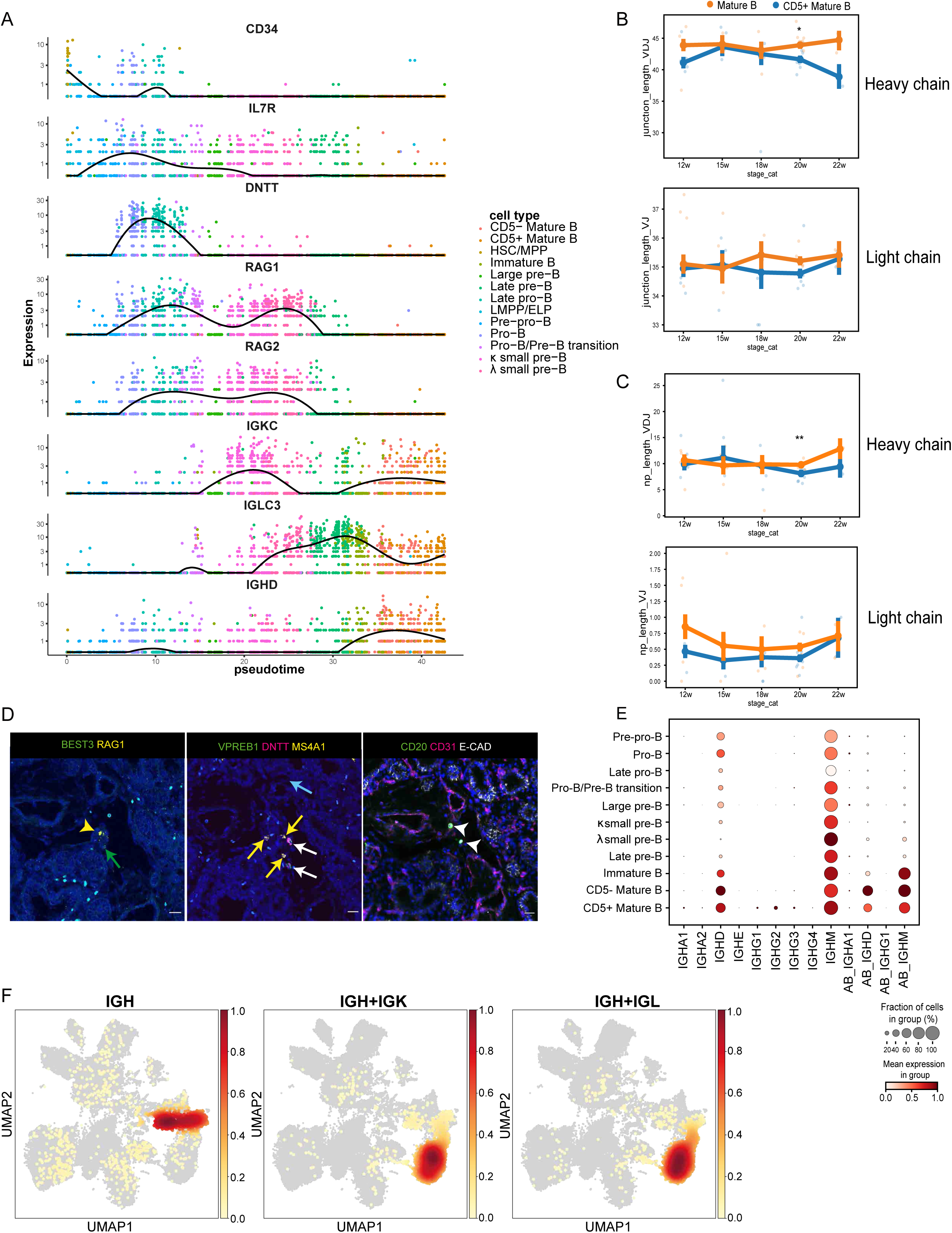
B cell developmental signatures. (**A**) Selection of differentially expressed genes using a spatial autocorrelation test over the principal graph in monocle3. The displayed genes show temporally distinct gene expression patterns over pseudotime. (**B**) CDR3 junction and (**C**) N/P insertion lengths (nucleotide) in the heavy chain (VDJ) and light chain (VJ) across timepoints. Lines indicate the mean ± SEM of lengths, using single cell values. Scatter points indicate sample mean values. CD5+ Mature B and CD5-Mature B points and lines are indicated as blue and orange respectively. Benjamini-Hochberg corrected pairwise wilcoxon rank sum test where *=p<0.05 and **=p<0.01 between cell types at each time point (using sample mean values). RNAscope using 18 pcw fetal lung sequential tissue sections (**D**, left and center) shows B cells expressing multiple combinations of markers, *BEST3*, *RAG1*, *VPREB1*, *DNTT* and *MS4A1* (yellow arrowheads (pre-B): *BEST3^+^RAG1^+^*, green arrows (large pre-B): *BEST3^+^*, white arrows (pro-B): *VPREB1^+^DNTT^+^*, blue arrows (large pre-B): *VPREB1^+^*, yellow arrows (late pre-B): *VPREB1^+^MS4A1^+^*). Corresponding IHC (**D**, right) using the next sequential tissue section, shows expression of CD20, CD31 (endothelium/blood vessels) and ECAD (epithelium) (arrowheads show location of CD20^+^ B cells). (**E**) shows *IGH* (immunoglobulin Heavy locus) gene expression in B cell populations from RNA+CITE-seq samples (‘AB_’ for protein measurements). (**F**) UMAPs show the density of immune receptor classes *IGH*, *IGK* (immunoglobulin Kappa locus) and *IGL* (immunoglobulin Lambda locus) in the single cell dataset. Related to Fig 3.

**Figure S5.**
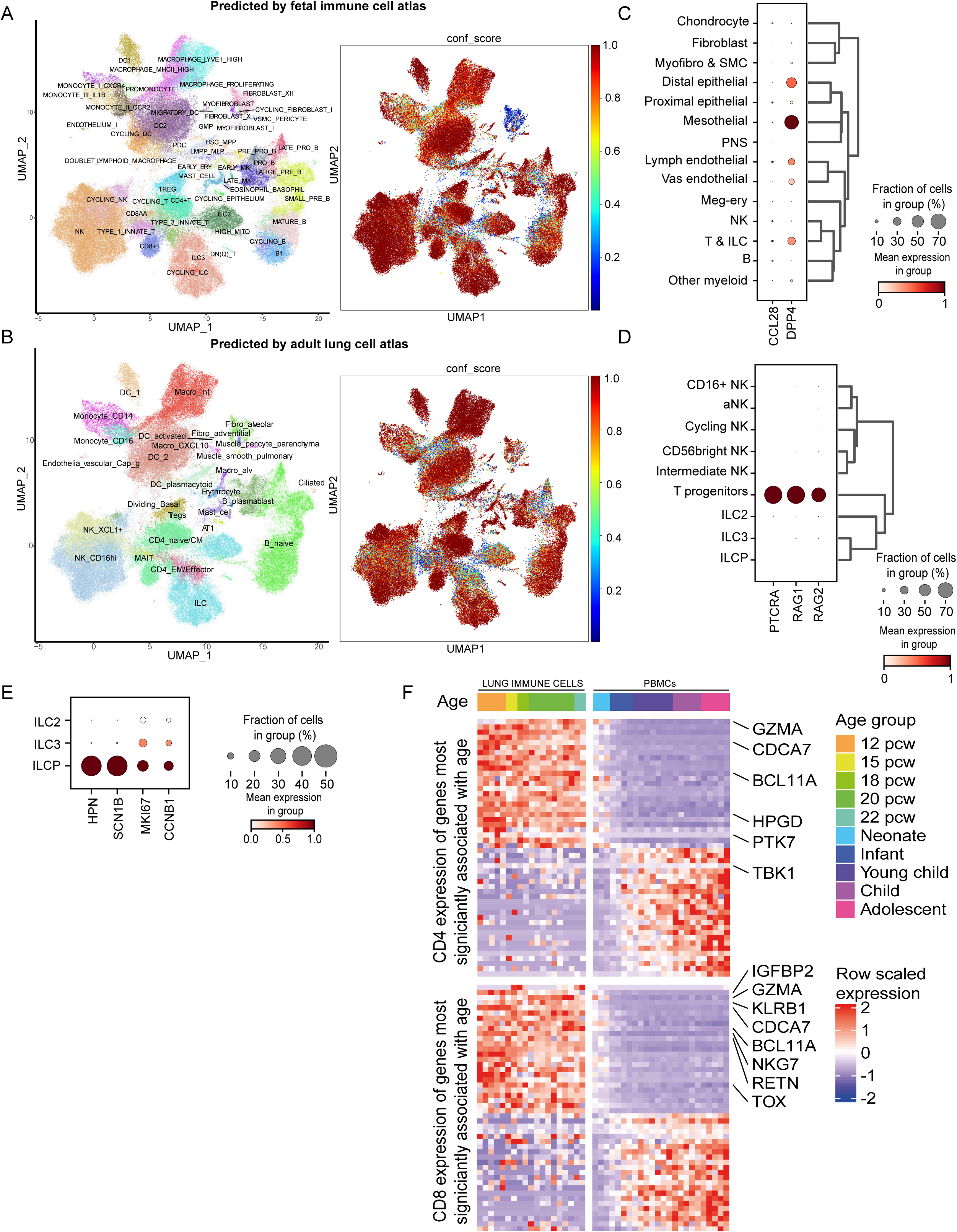
Comparison of fetal lung immune cells with published data. (**A,B**) CellTypist-predicted labels based on models trained from the pan-fetal immune data^20^ (**A**) and adult lung dataset^83^ (**B**). (**C**) Putative interacting chemokine-receptor partners enriched in B-1 cells. Marker genes in T progenitors (**E**) and ILCPs (**F**). (**G**) Relative expression dynamics of genes that are most significantly associated with age in naive CD4^+^ (top) and CD8^+^ (bottom) T cells. Genes reported to be related to T cell identity are highlighted. Postnatal (PBMC) data from healthy neonates and pediatric donors are derived from^35^. Related to Fig 4.

**Figure S6.**
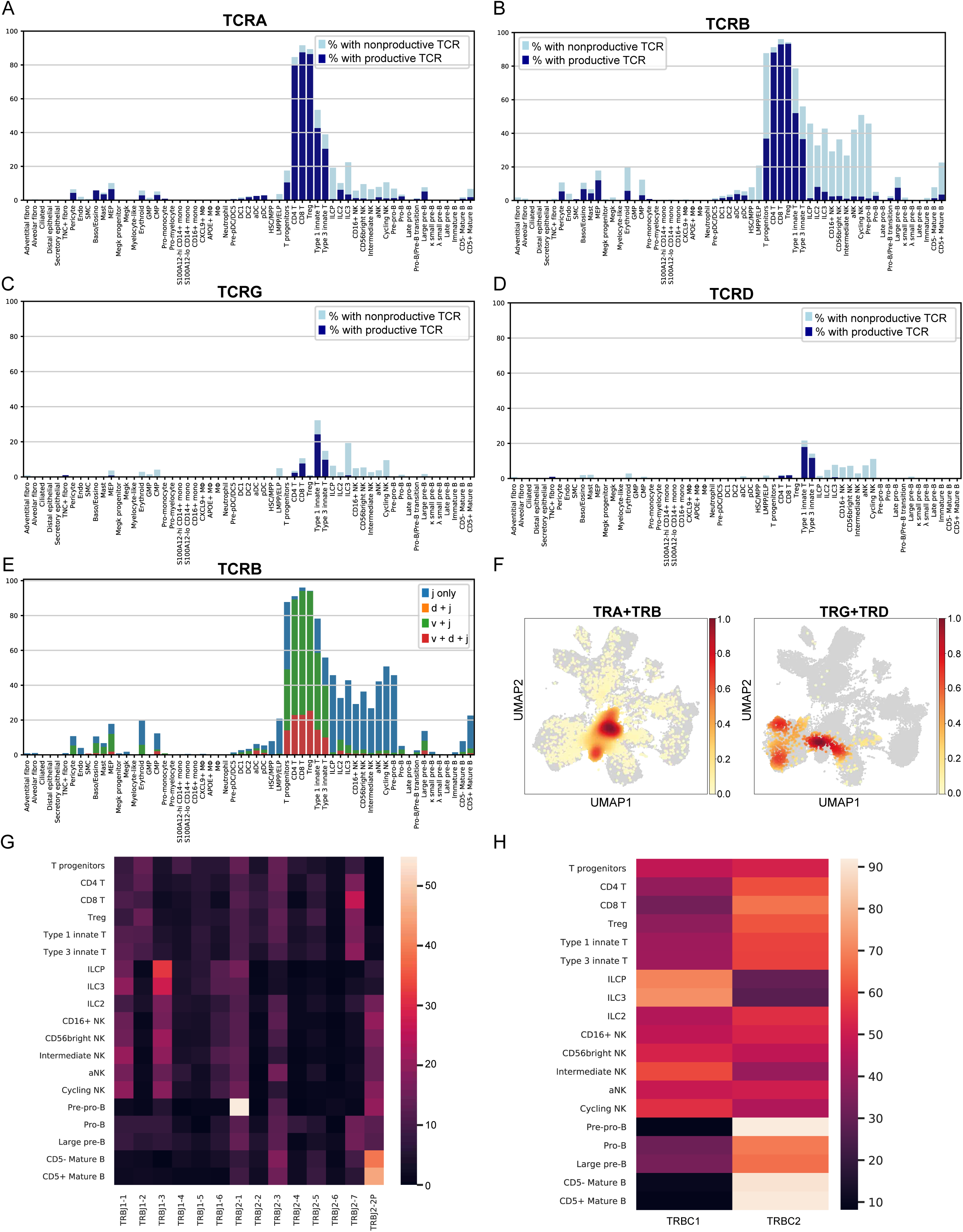
TCR Analysis. (**A**) Proportions of cells expressing productive or nonproductive TCRA contig. (**B**) Proportions of cells expressing productive or nonproductive TCRB contig. For (**A**) and (**B**), the proportions were calculated over cells that had single-cell abTCR sequencing. (**C**) Proportions of cells expressing productive or nonproductive TCRG contig. (**D**) Proportions of cells expressing productive or nonproductive TCRD contig. For (**C**) and (**D**), the proportions were calculated over cells that had single-cell gdTCR sequencing. (**E**) Proportions of cells expressing productive TCRB contig mapped to J chain only (j only), or D chain and J chain (d+j), or V chain and J chain (v+j), or V chain and D chain and J chain (v+d+j). The proportions were calculated over cells that had single-cell abTCR sequencing. (**F**) UMAPs showing the density of the T immune receptor classes within the single cell dataset. Heatmaps showing the percentage of each TRBJ gene segment present (**G**) and each TRBC gene segment present in different cell types (**H**). Related to Fig 4.

**Figure S7.**
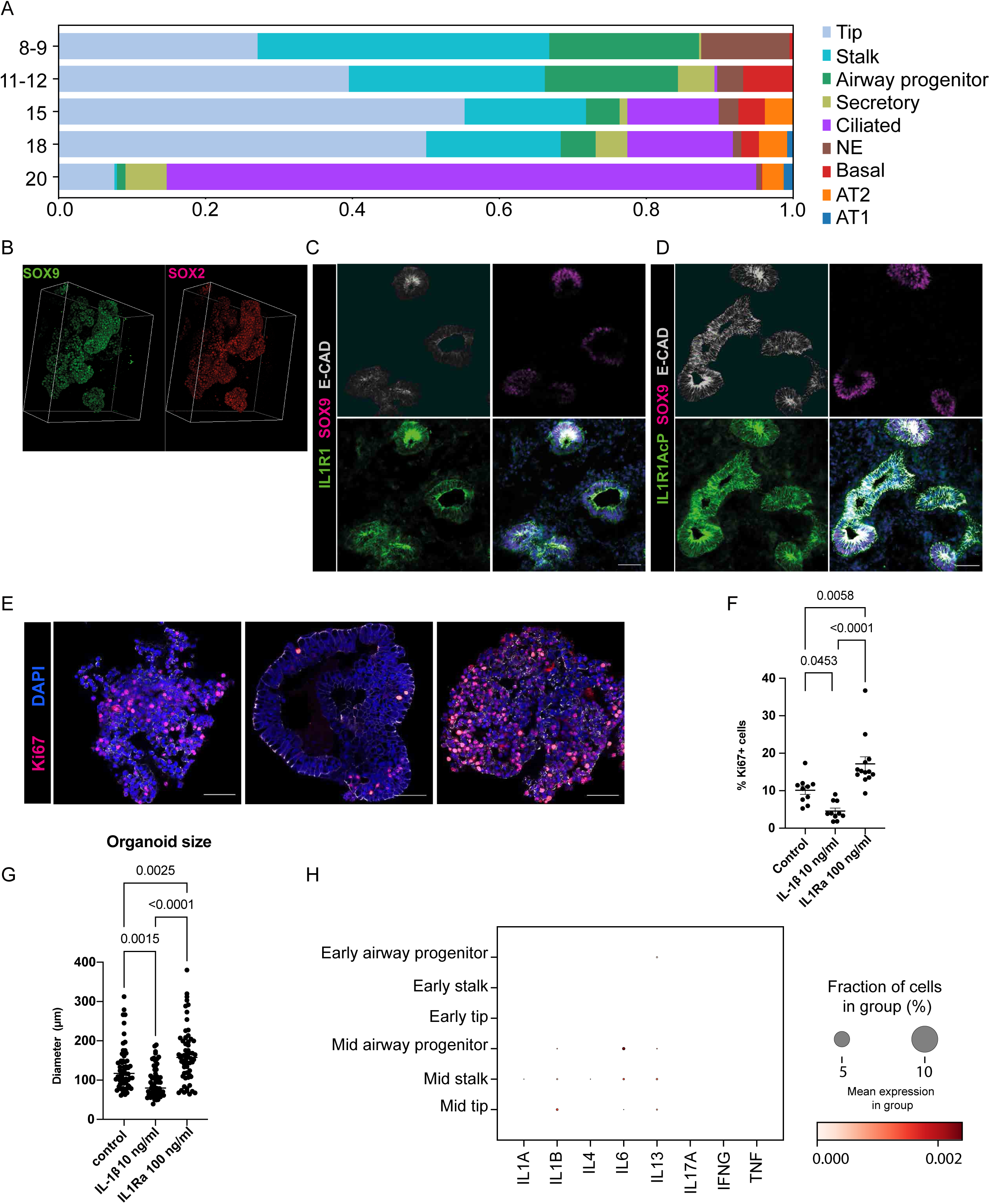
Additional organoid data. (**A**) Proportions of fetal lung broad epithelial cell types across different age groups, based on our previous single cell dataset^21^. (**B**) Whole mount 3D organoid staining shows stem cell markers, SOX2 and SOX9 (scale bar = 50μm). Representative IHC images show IL-1R1 (**C**) and IL-1R1AcP (**D**) staining, relative to the epithelium (ECAD^+^) and SOX9^+^ epithelial tip progenitors in fetal lung (blue: DAPI^+^ nuclei; scale bar = 50 µM). (**E**) Organoids were treated with either 10 ng/ml IL-1β or 100 ng/ml IL-1Ra for 7 days and whole mount stained for Ki67 and ECAD (blue: DAPI^+^ nuclei; scale bar = 50 µM). (**F**) Quantification of Ki67^+^ cells per image. (**G**) Quantification of organoid size. All data are presented as mean ± SEM, n=3 biological replicates. p-values are calculated by one-way ANOVA followed by Tukey’s post-hoc test. (**H**) Absence of cytokine gene expression in the distal lung based on our previous single cell dataset^21^. Related to Fig 5.

**Figure S8.**
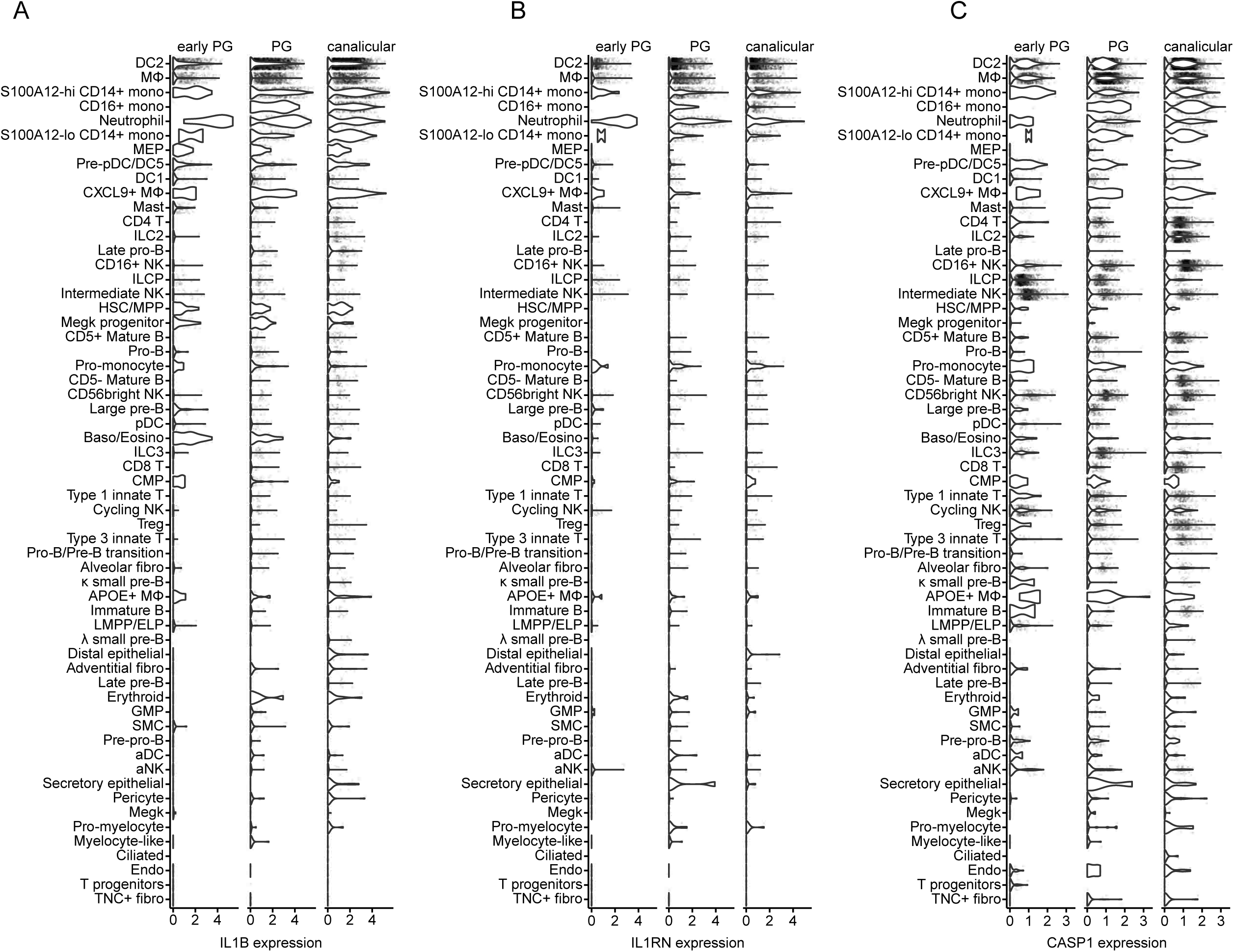
*IL1B*, *IL1RN* and *CASP1* expression and cell type prediction. Violin plots showing the cell type-specific expression of *IL1B* in (**A**), *IL1RN* in (**B**) and *CASP1* in (**C**). Analysis relates to Fig 6B, but shows expression in all cell types.

